# Skiver: reference-free quality control of metagenomic sequencing datasets using (*k, v*)-mer sketches

**DOI:** 10.64898/2026.02.12.705514

**Authors:** Zhenhao Gu, Puru Sharma, Limsoon Wong, Niranjan Nagarajan

## Abstract

Quality control is essential in numerous Bioinformatics pipelines, yet existing approaches often depend on inefficient mapping to reference genomes or potentially uncalibrated quality scores. We present *skiver*, a reference-free, alignment-free quality control tool that uses (*k, v*)-mer sketches. By deriving consensus from sketched (*k, v*)-mers, *skiver* estimates survival and hazard rates that capture the positional distribution of errors. Across simulated and real datasets from multiple sequencing platforms, *skiver* accurately estimates over-all error rates, error-type proportions, and quality-score calibration. An iterative outlier-filtering strategy enables robust performance on complex datasets with multiple strains, alleles, and repetitive regions.

## 1 Background

Given a set of sequenced reads, accurately estimating the overall sequencing error rate and the spectrum of error types (substitutions, insertions, and deletions) is a fundamental task in computational biology. These statistics are central to routine quality control (QC). They are used to decide whether a run is usable, to compare runs across different flow cells, chemistries, or library preparations, and to detect systematic failure modes [1]. They also directly affect downstream inference. In particular, error profiles influence alignment scoring and chaining heuristics in read mapping [2], and serve as key priors or likelihood components in sensitive variant calling [3] and strain-aware analysis such as phasing [4]. Because many pipelines implicitly assume a particular error model, biased error rate estimates can propagate and lead to avoidable miscalls, unstable parameter tuning, and misleading biological conclusions.

This challenge is amplified in metagenomic settings. Real samples often contain a mixture of organisms with highly uneven abundance, including low-coverage genomes and closely related co-existing strains. These properties can confound error rate estimation in two ways: (i) low-coverage genomes and strain heterogeneity lead to many *k*-mers with low coverage, which can be mistaken for sequencing errors, and (ii) reference genomes may be incomplete or unavailable because organisms are missing from databases or differ substantially from available references. Consequently, methods that rely on an incomplete reference genome database can produce biased estimates precisely in the scenarios where robust error profiling is most needed.

### 1.1 Previous work

Existing approaches for estimating sequencing error rates and spectra can be broadly divided into reference-based and reference-free methods. Reference-based methods infer errors from alignments to an external reference, whereas reference-free methods avoid mapping and instead rely on internal read statistics such as *k*-mer frequency patterns or basecalling quality scores. Below, we summarize the limitations of each category, with an emphasis on failure modes that are especially pronounced in metagenomic samples.

#### Reference-based methods

A common and often effective strategy is to map the reads to a reference genome and compute error statistics from the resulting alignments (e.g., via CIGAR operations) [2, 5]. When a high-quality, closely matching reference is available, this approach can provide accurate estimates. However, it has two major drawbacks. First, mapping and alignment can be computationally expensive for large read sets and for long references, making it costly as a routine QC step. Second, and more critically for metagenomics, suitable references may be missing, incomplete, or diverge from the sequenced genomes because organisms are not represented in databases or because strains differ substantially from available references. In such cases, true biological differences are conflated with sequencing errors, inflating apparent mismatch and indel rates and biasing the estimated sequencing error rates upward [6]. One possible mitigation is to assemble the reads to obtain a sample-specific consensus and then compare the reads to this consensus [7], but assembly can be computationally intensive and may be unstable for low-coverage components and complex mixtures.

#### Reference-free methods

To avoid dependence on an external reference, many approaches estimate error rates directly from read-derived statistics. For example, *shadow regression* [8] and *SequencErr* [9] utilize overlapping (paired-end) short reads to quantify disagreements between reads. Another prominent family of methods lever-ages *k*-mer frequency information to infer error-related quantities without explicit alignment [10–17]. While scalable, these methods are often tailored to specific regimes (e.g., single-genome assumptions or specific sequencing platforms) and can struggle to recover the full error spectrum (i.e., distribution of substitutions vs. insertions vs. deletions). Moreover, the severe coverage imbalance typical of metagenomic samples introduces a key pitfall: low-coverage genomes yield sparse and highly variable *k*-mer counts, making frequency-based inference unstable and causing rare error-free *k*-mers to be difficult to distinguish from erroneous *k*-mers induced by sequencing noise.

Another class of reference-free methods uses Phred quality scores, which theoretically satisfy *Q* = −10 log_10_ Pr[error] and are used by many downstream tools [3, 18]. However, in practice, quality scores can be miscalibrated and vary with sequencing technology [19] and library preparation [20]. Some calibration procedures do exist [21], but require additional assumptions and data. Consistent with these observations, our experiments show that uncalibrated quality scores can substantially underestimate or overestimate true error rates. Moreover, they do not directly provide reliable estimates of the full substitution/insertion/deletion spectrum.

### 1.2 Our contribution

We propose *skiver*, which introduces a novel (*k, v*)-mer sketching approach for reference-free and alignment-free profiling of sequencing error rates and spectra. The core idea is to construct sketches that group (*k, v*)-mers sharing the same *key* and then find the error patterns in the associated *values*. Using concepts in survival analysis, we use the hazard rate and the survival rate to describe the frequencies and the distribution of sequencing errors, and model the number of bases until the first sequencing error using a discrete Weibull distribution. Across simulated and real datasets from multiple sequencing platforms, our method yields accurate hazard rate and survival rate estimates while remaining robust in settings where reference genomes are unavailable or incomplete, including metagenomic regimes with uneven coverage and strain heterogeneity.

### 1.3 Problem formulation

There are various definitions of sequencing error rate in the literature, including the BLAST error rate, gap-compressed error rate, and gap-excluded error rate [22]. Among these, the BLAST error rate is the most commonly used and is defined as the number of errors divided by the alignment length [23]. In this paper, we borrow concepts from survival analysis. Let *T* (*T* ≥1) be the random variable that is the number of bases from a random starting position until the first failure (disagreement between the sequenced base and the true underlying base).

#### Definition 1

The *hazard rate* at the *t*-th base from the starting position, denoted by *h*(*t*) := Pr[*T* = *t* | *T* ≥ *t*], is the conditional probability of the *t*-th base disagreeing with the underlying genome given that the previous (*t* − 1) bases agree.

The benefits of using the hazard rate instead of the error rate inferred from alignment include the following. Firstly, alignment can be slow and ambiguous. Different alignment scoring schemes may yield different numbers of aligned bases, resulting in varying estimates using different mappers [24]. Secondly, the hazard rate offers a more comprehensive view of the position-dependent error process. For example, imagine two reads in **Figure 1**, where read A has 4 consecutive errors every 16 bases, and read B has 1 error every 4 bases. The error rates of the two reads are the same and are equal to 1/4, but their hazard rates are drastically different. Specifically, read A has a much lower hazard rate at small *t*, indicating a higher chance of observing a long stretch of consecutive matches.

**Fig. 1.**
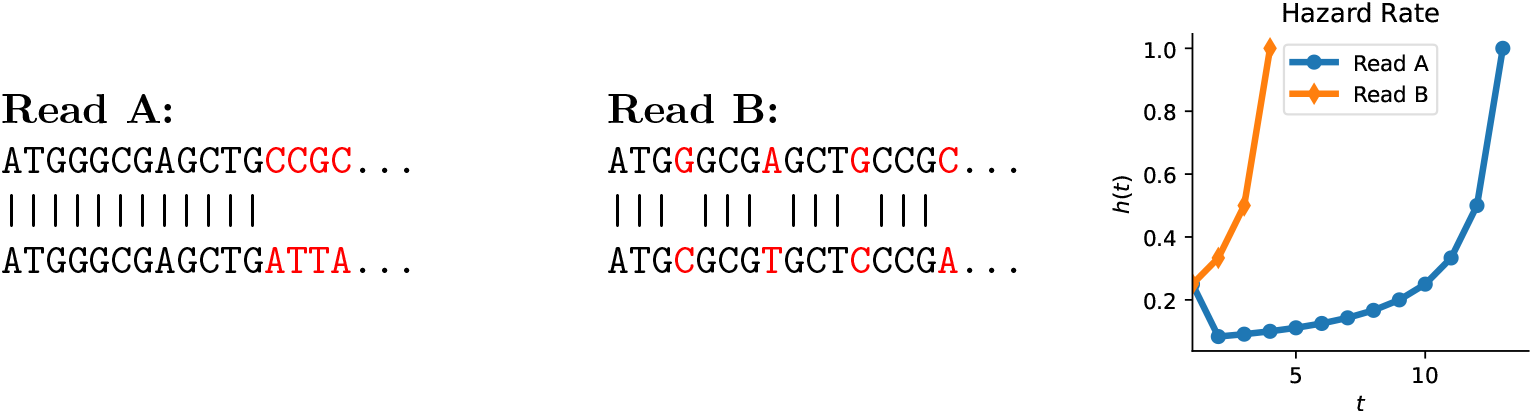
The alignment of two reads (bottom sequence) to the reference (top sequence). Both reads have an error rate of 1/4 but different hazard rates *h*(*t*) (right).

#### Definition 2

The *survival rate* at the *t*-th base, denoted *S*(*t*) := Pr[*T > t*], is the probability of the first *t* bases from the point of observation being free of sequencing errors.

Note that the survival rate 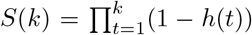 also indicates the probability of a *k*-mer in the read being free of sequencing error. The *per-base error rate* can be calculated by *ε*_per-base_ := *h*(1) = 1 −*S*(1), which is the probability that a random base in the read is erroneous.

In this work, we propose an efficient and accurate estimator for hazard rate and survival rate using (*k, v*)-mer sketches, and show its ability to estimate the frequency of each error type (substitutions, insertions, and deletions) at the same time.

## 2 Results

### 2.1 Baselines and datasets

To test the ability of our algorithm to estimate *h*(*t*) and *S*(*t*), we benchmarked skiver v0.2.2 against popular state-of-the-art baselines.

For mapping-based tools, we used BEST [5] to analyze the BAM output from Minimap2 [2], with ground truth reference genomes. By default, BEST uses only the primary alignment for each read for read error profiling. For reference-free methods, we used GenomeScope2.0 [16] to fit the *k*-mer histogram output of KMC 3.2.4 [29], and used the `Read Error Rate` field in the summary file. For quality-score-based methods, we used seqtk [30] and the `ErrQ` field to compute sequencing error rates,

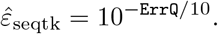

We selected a wide range of sequencing datasets from multiple sources and sequencing platforms (**Table 1**) [25–28, 31], including mock bacterial communities, bacterial isolates, human reads, and different microbiomes. All of the chosen datasets (except the microbiomes) have well-defined reference genomes for fair comparisons.

**Table 1.**
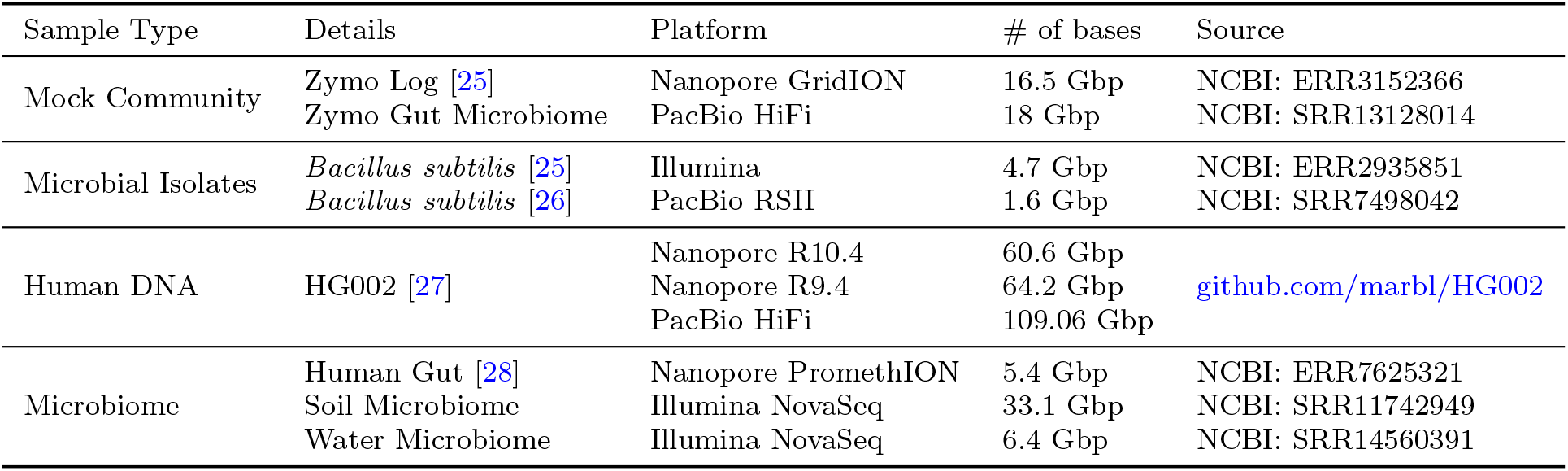
Sequencing datasets used for various experiments in this study.

| Sample Type | Details | Platform | # of bases | Source |
|---|---|---|---|---|
| Mock Community | Zymo Log [25] | Nanopore GridION | 16.5 Gbp | NCBI: ERR3152366 |
|  | Zymo Gut Microbiome | PacBio HiFi | 18 Gbp | NCBI: SRR13128014 |
| Microbial Isolates | *Bacillus subtilis* [25] | Illumina | 4.7 Gbp | NCBI: ERR2935851 |
|  | *Bacillus subtilis* [26] | PacBio RSII | 1.6 Gbp | NCBI: SRR7498042 |
| Human DNA | HG002 [27] | Nanopore R10.4 | 60.6 Gbp | github.com/marbl/HG002 |
|  |  | Nanopore R9.4 | 64.2 Gbp |  |
|  |  | PacBio HiFi | 109.06 Gbp |  |
| Microbiome | Human Gut [28] | Nanopore PromethION | 5.4 Gbp | NCBI: ERR7625321 |
|  | Soil Microbiome | Illumina NovaSeq | 33.1 Gbp | NCBI: SRR11742949 |
|  | Water Microbiome | Illumina NovaSeq | 6.4 Gbp | NCBI: SRR14560391 |

All experiments were run on a server with 16 Intel(R) Xeon(R) Platinum 8259CL CPUs @ 2.50GHz. All baseline tools were run with their default parameters.

### 2.2 Evaluation metrics

In all of the experiments, we calculate the ground truth survival rate *S*(*t*) by counting the proportion of *t*-mers in the reads that align with the ground truth reference without any error in the BAM output of Minimap2. The ground truth hazard rate was calculated as *h*(*t*) = 1 −*S*(*t*)*/S*(*t* −1) for *t* ≥2 and *h*(1) = 1 −*S*(1).

The main metric used to evaluate the tools was the mean squared error (MSE) of the estimated survival rate *Ŝ*(*t*) vs. the ground truth *S*(*t*), calculated by

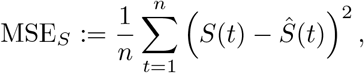

where the upper bound *n* is chosen to be 100. This is similar to the definition of Brier scores, which are used extensively in the evaluation of survival curve estimation [32]. Intuitively, this metric measures how well the tools predict the proportion of the *t*-mers in the read set that are error-free for various *t*. Similarly, we define the MSE for hazard rate to be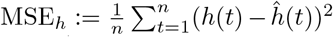. Here, 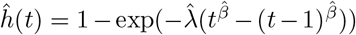 and 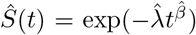 for skiver, while 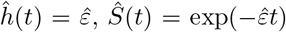 for the other methods that only report one error rate estimate 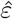.

### 2.3 Results for mapping-based methods may be biased by wrong or missing references

In this section, we evaluate the behavior of mapping-based methods under simulated conditions in which the reference genome is missing or incorrect.

We first consider the scenario of an incomplete reference database, a common situation in metagenomic analyses. Using the *Zymo Log, Nanopore GridION* dataset, we applied Minimap2 to map the reads under two reference configurations: (i) using all 10 known genomes in the mock community as references, and (ii) using each individual genome alone as the reference (**Figure 2.A**).

**Fig. 2.**
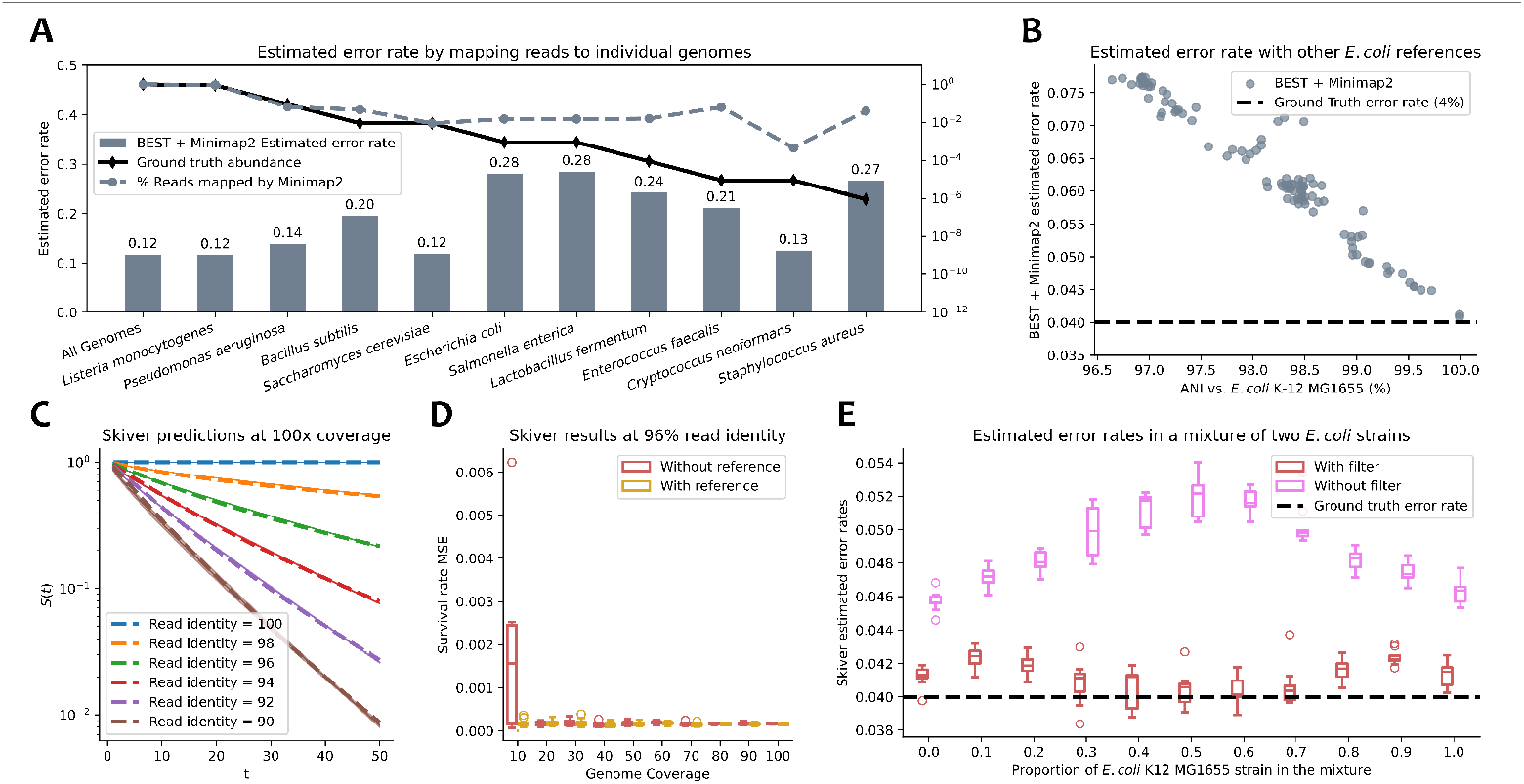
Performance of various tools under simulated conditions. **A**. Profiled error rate for BEST + Minimap2 on the Zymo Log mock community dataset, if only one bacterial isolate is given as the reference (on the x-axis). The lines representing the abundance and the % reads mapped use the axis on the right of the figure. **B**. Profiled error rate for BEST + Minimap2 for a simulated read from an *E. coli* K-12 MG1655 strain, using different *E. coli* strains as references. **C**. Estimation of survival rates *S*(*t*) for simulated datasets of *E. coli* with 100 *×* coverage. The dashed line represents the ground truth *S*(*t*) at different read identities (%), the solid line represents the mean of the estimates *Ŝ*(*t*), and the shaded area covers the range between the min and max of *Ŝ*(*t*) in the 20 simulated experiments. **D**. MSE_*S*_ of skiver estimates at different coverage values with simulated reads that have 96% read identity. **E**. Estimated error rates for skiver when mixing two *E. coli* strains together at different proportions, with and without the outlier filter.

Interestingly, the estimated error rate obtained using the complete reference set is the lowest (approximately 12%), whereas estimates derived from individual reference genomes are consistently biased upwards. Examination of the mapping results reveals that Minimap2 assigns a disproportionately large number of reads to certain isolate genomes relative to their expected abundances in the mock community. This indicates that many reads originating from other species are incorrectly mapped to these genomes. As a result, the apparent sequencing error rate is substantially inflated. These observations suggest that read mapping with an incomplete reference database is inherently unreliable for error-rate profiling. While it may be possible to reduce spurious mappings by applying more stringent alignment score thresholds, the optimal threshold itself depends on the unknown sequencing error rate, creating a circular dependency that limits the robustness of mapping-based approaches.

Next, we examine the case in which the reference genome is present but does not exactly match the sequenced strain. Reads were simulated from *E. coli* K-12 MG1655 at 96% read identity using Badread [33]. Error rates were then estimated using Minimap2 and BEST, with reference genomes chosen from other *E. coli* strains sampled from GTDB [34]. The average nucleotide identity (ANI) between each reference genome and *E. coli* K-12 MG1655 was computed using skani [35], and all selected references were verified to have ANI values greater than 95%. Despite this high sequence similarity, the estimated error rate almost doubled as the ANI between the reference genome and the true sequenced genome decreased to 96% (**Figure 2.B**). These results demonstrate that without an additional step of variant calling, even modest strain-level divergence can severely bias error-rate estimates derived from read mapping. Consequently, accurate profiling using mapping-based methods requires the exact strain, or a near-identical genome, to be present in the reference database, a condition that is rarely met in real metagenomic samples.

### 2.4 Skiver accurately estimates hazard rates and survival rates in simulated reads

Next, we tested the behavior of skiver on simulated reads.

We first evaluated skiver under a simplified setting in which the read set contains only a single genome. Reads from the *E. coli* K-12 MG1655 strain were simulated across a range of read identities (90% to 100%) and sequencing coverages (10×to 100×). We performed 20 experiments on each set of parameters.

Across these conditions, skiver accurately recovered both the survival rate and the hazard rate. For example, at 100 ×coverage (**Figure 2.C**), the estimated survival curve *Ŝ*(*t*) (solid line) closely matches the ground-truth *S*(*t*) (dashed line), exhibiting minimal bias and low variance. When the read identity is fixed (**Figure 2.D**), the mean squared error of the survival rate estimate (MSE_*S*_) remains close to zero as coverage decreases, until coverage drops to 10×. At this depth, the default skiver configuration shows increased variance, whereas providing a reference genome restores accuracy and reduces variance.

Based on the complete set of simulations (**Supplementary Figures S1-2**), skiver produces generally reliable survival and hazard rate estimates when at least one genome in the read set has coverage exceeding 20×, and the read error rate is below 6%. For lower coverage or higher error rates, a reference genome is required for an accurate estimate.

In addition to accurately estimating survival and hazard functions, skiver is also able to recover the underlying sequencing error spectrum used by the simulator. In this experiment, Badread was configured with a random error model, in which both the type of edit operation (substitution, insertion, or deletion) and the resulting base after each edit are selected uniformly at random. Consistent with this design, the error spectrum reported by skiver (**Figure 3**) exhibits approximately equal frequencies across all single-base substitution (SBS) types, while the overall frequencies of substitutions, insertions, and deletions are each close to one third. These results closely match the parameters specified in the simulator.

**Fig. 3.**
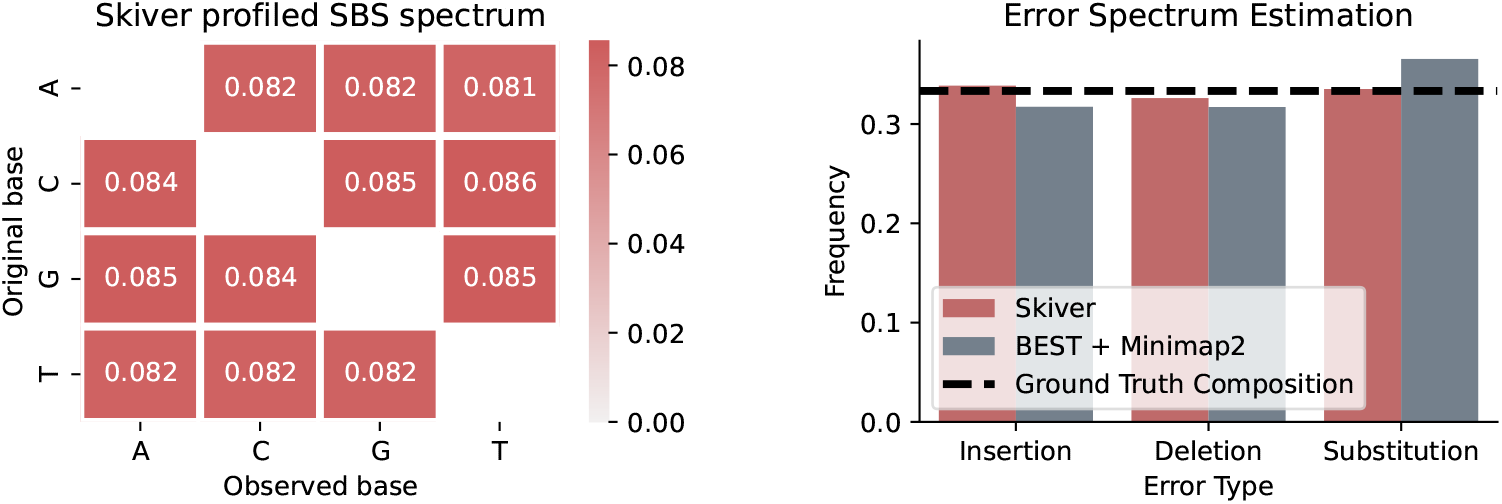
Estimated single-base substitution (SBS) spectrum of a 128× coverage *E. coli* dataset with 96% read identity simulated using Badread, where all error types are equally likely.

In contrast, when BEST is applied to Minimap2 alignments generated from the same dataset, a higher proportion of substitutions is reported relative to insertions and deletions. This bias is likely attributable to the alignment scoring scheme of Minimap2, which favors mismatches over indels during alignment optimization. By comparison, skiver is an alignment-free method and therefore remains unaffected by alignment heuristics or scoring parameters, enabling a more faithful recovery of the true sequencing error spectra.

Next, we evaluated the effect of the outlier filter in datasets containing multiple closely related strains (**Figure 2.E**). We simulated reads with 64× coverage from *E. coli* K-12 MG1655 and *E. coli* O157:H7 at the same read identity. These two strains share an average nucleotide identity (ANI) of 98% according to skani [35]. The two read sets were then subsampled and mixed to generate datasets with a fixed total coverage of 64*×* but varying proportions of the two strains. For each mixture, we estimated the sequencing error rate using skiver, defined as the first-base hazard 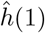, both with and without the outlier filter enabled.

When the proportion of K-12 MG1655 is 1.0 or 0.0, the dataset effectively represents a single isolate. In these cases, skiver without filtering reports overestimated error rates, largely due to similar genomic regions that introduce spurious (*k, v*)-mer groupings. As the mixture proportion approaches 0.5, the estimated error rate from skiver without filtering increases sharply, reflecting the growing ambiguity between orthologous regions shared by the two strains. In contrast, skiver with the outlier filter remains relatively robust across all mixture proportions and consistently reports error rates close to the ground truth. These results demonstrate that the outlier filter effectively suppresses confounding signals arising from repetitive and shared regions in strain genomes, enabling accurate error-rate estimation even in multi-strain datasets.

### 2.5 Skiver generalizes well to reads from various sequencing platforms

We next evaluated the performance of skiver and several baseline methods on real sequencing datasets.

Across all datasets, skiver produces accurate hazard rate estimates (**Figure 4.A**). For Illumina reads, the estimated hazard rates are slightly biased upward, which is likely attributable to known positional biases in Illumina sequencing, where errors are more prevalent at the two ends of the reads (**Supplementary Figure S3; Supplementary Table S1**). Notably, the estimated 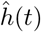 curves are nearly identical for skiver with and without a reference genome, indicating that selecting the most frequently observed value as the consensus is a reliable strategy.

**Fig. 4.**
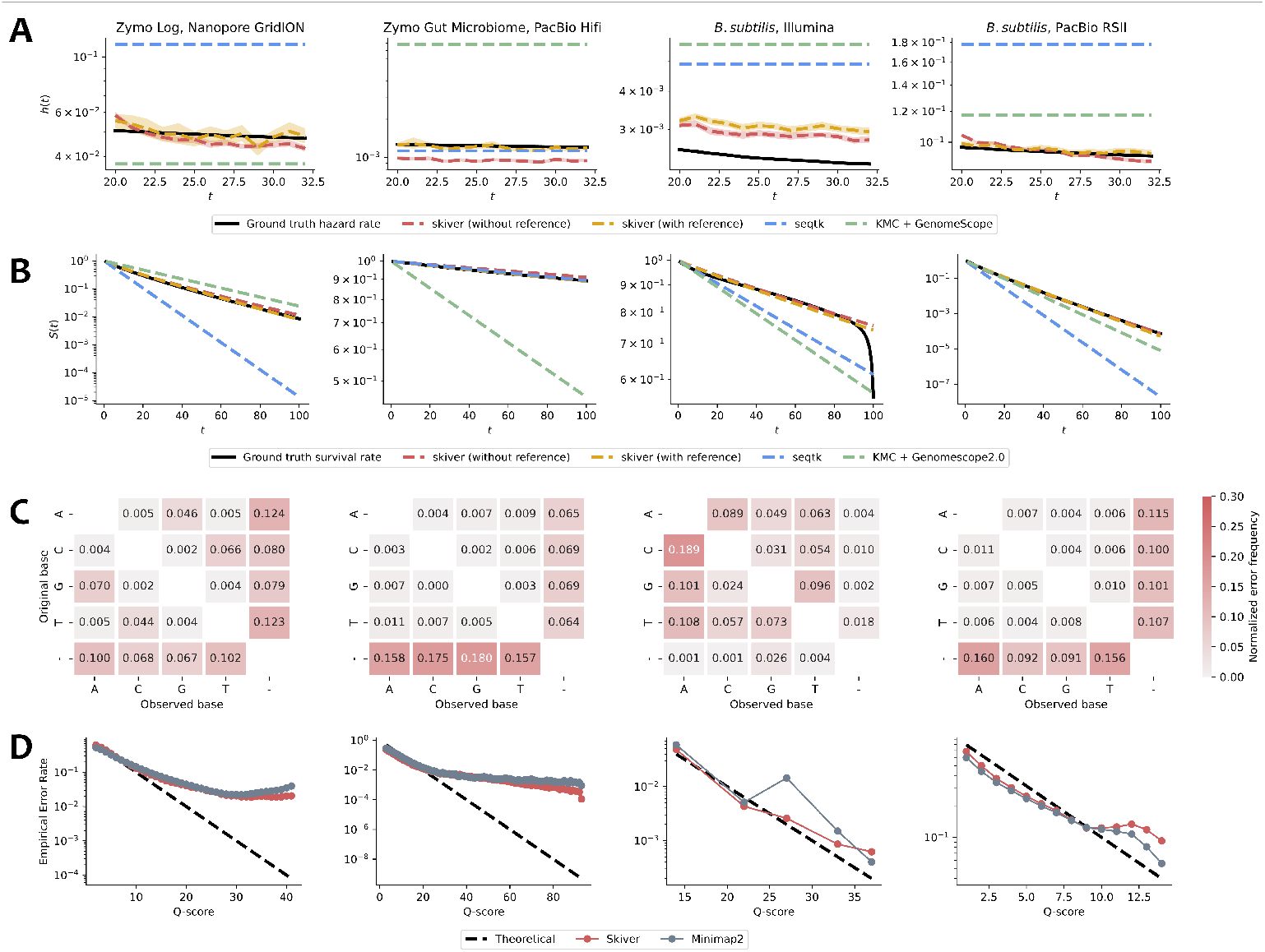
Performance of skiver on real metagenomic datasets. **A**. Estimated hazard rate using skiver, with and without reference genomes. The shaded areas are the 90% confidence intervals for the hazard rate estimated via bootstrapping. **B**. Estimated survival rate using baselines and skiver. **C**. Estimated error spectra for the datasets. The last row represents insertions, and the last column represents deletions, while the rest of the entries are substitutions. **D**. Estimated true error rate of the bases with respect to their associated Phred scores.

For survival rate estimation, skiver demonstrates consistent performance, achieving low MSE_*S*_ and MSE_*h*_ across all datasets (**Figure 4.B; Table 2; Supplementary Table S2**). Skiver also produces a reasonable estimate of error rates across multiple real complex metagenomic samples that agrees with previous error profiles (**Supplementary Table S3**). Among the baselines, KMC + GenomeScope 2.0 is not designed for metagenomic samples and therefore struggles to identify an appropriate cutoff between erroneous and solid *k*-mers in the *k*-mer count histogram. Its performance is nevertheless substantially better on bacterial isolates and human sequencing data. Seqtk, on the other hand, frequently overestimates error rates, reflecting the fact that base quality scores in many read sets are not well calibrated.

**Table 2.** MSE_*S*_ for benchmarked tools on real sequencing datasets. The tool with the lowest MSE_*S*_ is marked in bold, and the second lowest is marked with an underline.

| Dataset | seqtk | KMC + GenomeScope2.0 | skiver | skiver (with reference) |
|---|---|---|---|---|
| Zymo Log, Nanopore GridION | 1.28 × 10<sup>−2</sup> | 1.05 × 10<sup>−2</sup> | **1.47 × 10<sup>−4</sup>** | 1.78 × 10<sup>−4</sup> |
| Zymo Gut Microbiome, PacBio HiFi | 1.77 × 10<sup>−5</sup> | 7.95 × 10<sup>−2</sup> | 1.74 × 10<sup>−4</sup> | **1.56 × 10<sup>−6</sup>** |
| *B. subtilis*, Illumina | 6.56 × 10<sup>−3</sup> | 1.24 × 10<sup>−2</sup> | 6.80 × 10<sup>−4</sup> | **5.65 × 10<sup>−4</sup>** |
| *B. subtilis*, PacBio RSII | 5.60 × 10<sup>−3</sup> | 3.61 × 10<sup>−4</sup> | 4.71 × 10<sup>−4</sup> | **6.16 × 10<sup>−5</sup>** |
| Human, Nanopore R10.4 | 2.35 × 10<sup>−1</sup> | 1.47 × 10<sup>−2</sup> | 1.63 × 10<sup>−3</sup> | **6.33 × 10<sup>−4</sup>** |
| Human, Nanopore R9.4 | 8.60 × 10<sup>−2</sup> | 5.44 × 10<sup>−2</sup> | **1.58 × 10<sup>−3</sup>** | 3.06 × 10<sup>−3</sup> |
| Human, PacBio HiFi | 7.93 × 10<sup>−3</sup> | 1.70 × 10<sup>−4</sup> | 1.78 × 10<sup>−4</sup> | **5.51 × 10<sup>−6</sup>** |

Since skiver accurately estimates the survival rate *S*(*k*), i.e., the probability that a *k*-mer is free of sequencing errors, its output can serve as prior knowledge for downstream *k*-mer-based methods. For example, skiver’s error rate estimation can be used as input to sylph [36], a fast metagenomic profiling tool, to accurately estimate the percentage of unclassified reads (**Supplementary Table S4**).

In addition to estimating error rates, skiver enables profiling of sequencing error spectra (**Figure 4.C**). For nanopore datasets, substitutions are dominated by `G→A` and `A →G` transitions; PacBio datasets are characterized primarily by insertions and deletions; and Illumina datasets are dominated by substitution errors. These patterns are consistent with previously reported error profiles for the respective sequencing platforms [19, 24, 37]. The estimated spectra produced by skiver are also highly consistent regardless of whether a reference genome is provided (**Supplementary Figure S4**), further supporting the idea that the consensus derived from the most frequently observed value recovers the true reference base with high probability. On the other hand, the error spectra estimated using BEST in combination with Minimap2 differ noticeably from those obtained by skiver (**Supplementary Figure S5**). In particular, substitution rates are frequently inflated, which may be due to the alignment scoring scheme employed by Minimap2 and soft clipping.

Furthermore, skiver is able to calibrate Phred scores reported in the FASTQ files by estimating the conditional hazard rates, reporting similar empirical error rates as Minimap2 (**Figure 4.D**). In Minimap2, a base and its associated Phred score are considered as an error if it is a substitution, an insertion, or follows a deletion. In all cases, the empirical error rate decreases as the Phred score increases. Interestingly, across multiple sequencing platforms, the empirical error rate for low Phred scores (below 10) tends to be lower than the theoretical line, while high Phred scores are usually overconfident. The ability to calibrate Phred scores can potentially help downstream tools that rely on accurate Phred scores, such as LoFreq [3].

Finally, we compared the memory usage and running times of all evaluated tools (**Figure 5; Supplementary Tables S8-10**). Although skiver currently operates in a single-threaded mode, it remains among the fastest and most memory-efficient methods evaluated. This efficiency highlights the suitability of skiver as a lightweight and scalable first step in bioinformatics pipelines, particularly for large-scale analysis of sequencing datasets.

**Fig. 5.**
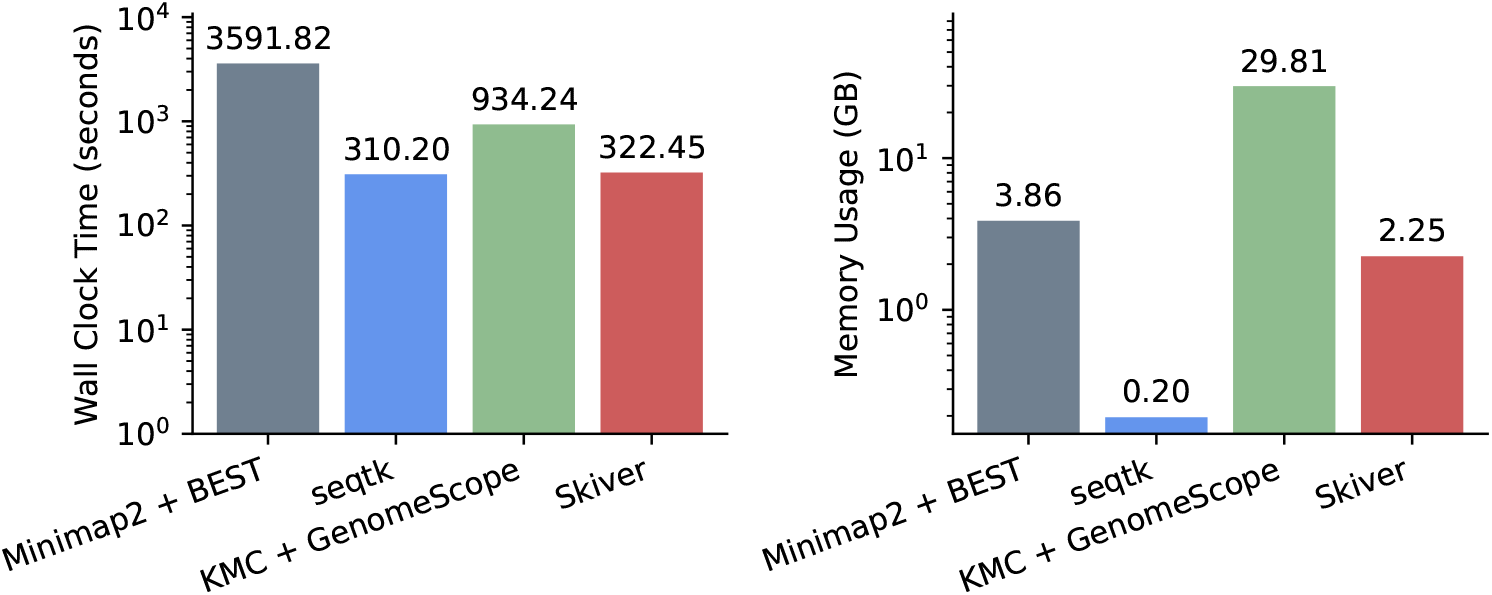
The wall clock time (left) and peak memory usage (right) of the profiling tools on the *Zymo Log, Nanopore GridION* dataset.

### 2.6 Ablation Studies

Lastly, we assessed whether modeling the time to first sequencing error, *T*, using a discrete Weibull distribution is appropriate. To this end, we constructed a variant of skiver in which the shape parameter is fixed at 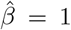. Under this constraint, the survival function simplifies to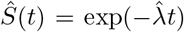, corresponding to a constant hazard rate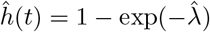.

Experiments conducted on the same datasets (**Table 3; Supplementary Table S7; Supplementary Figure S8**) show that this constant-hazard model still outperforms all baseline methods in terms of both MSE_*S*_ and MSE_*h*_ in most datasets, and provides a reasonable single-number estimate of the sequencing error rate. However, the full skiver model demonstrates a better average performance. The performance gap is particularly pronounced for Nanopore datasets, where sequencing errors are known to be non-uniform and tend to cluster along reads. These results indicate that the discrete Weibull distribution more closely reflects real sequencing error processes than a constant-hazard assumption. The difference can also be seen from the per-base error rate estimations, where the estimations of skiver with a constant hazard rate are much lower than the default skiver and Minimap2 + BEST’s estimations (**Supplementary Tables S3 and S6**).

**Table 3.**
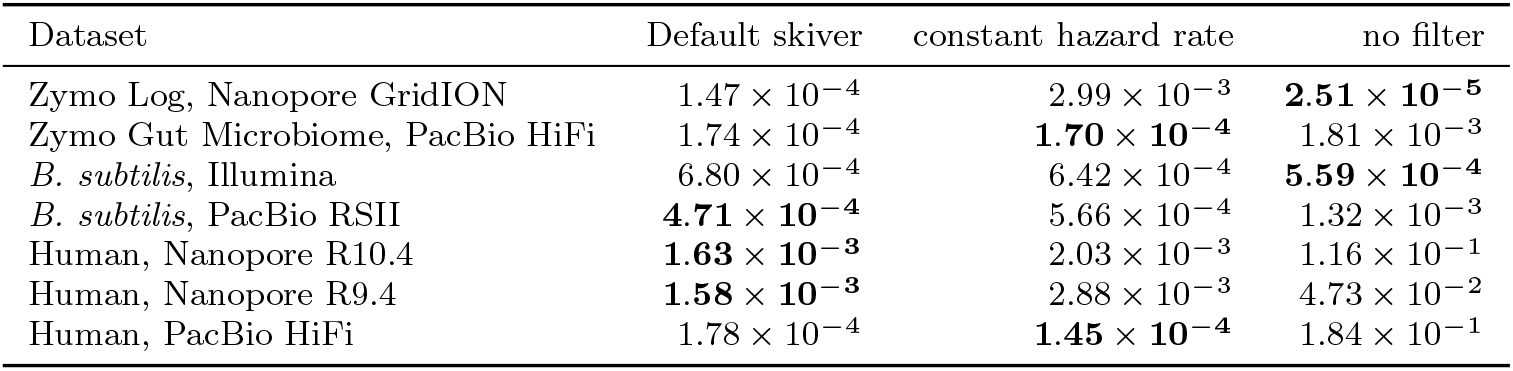
MSE_*S*_ across different datasets for variants of the skiver algorithm. “constant hazard rate”: if 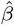is fixed to 1; “no filter”: if the outlier filter is disabled during hazard rate estimation. The lowest MSE_*S*_ is marked in bold.

| Dataset | Default skiver | constant hazard rate | no filter |
|---|---|---|---|
| Zymo Log, Nanopore GridION | 1.47 × 10<sup>−4</sup> | 2.99 × 10<sup>−3</sup> | **2.51 × 10<sup>−5</sup>** |
| Zymo Gut Microbiome, PacBio HiFi | 1.74 × 10<sup>−4</sup> | **1.70 × 10<sup>−4</sup>** | 1.81 × 10<sup>−3</sup> |
| *B. subtilis*, Illumina | 6.80 × 10<sup>−4</sup> | 6.42 × 10<sup>−4</sup> | **5.59 × 10<sup>−4</sup>** |
| *B. subtilis*, PacBio RSII | **4.71 × 10<sup>−4</sup>** | 5.66 × 10<sup>−4</sup> | 1.32 × 10<sup>−3</sup> |
| Human, Nanopore R10.4 | **1.63 × 10<sup>−3</sup>** | 2.03 × 10<sup>−3</sup> | 1.16 × 10<sup>−1</sup> |
| Human, Nanopore R9.4 | **1.58 × 10<sup>−3</sup>** | 2.88 × 10<sup>−3</sup> | 4.73 × 10<sup>−2</sup> |
| Human, PacBio HiFi | 1.78 × 10<sup>−4</sup> | **1.45 × 10<sup>−4</sup>** | 1.84 × 10<sup>−1</sup> |

We further performed the same ablation experiments using skiver with the outlier filter disabled. In this setting, the estimated survival rates are substantially lower (**Supplementary Figure S8**), particularly for the Zymo Gut Microbiome mock community, where multiple *E. coli* strains coexist, and for human datasets, which are diploid and contain significant amounts of repetitive regions. The improved agreement between the default skiver estimates and the ground truth in these challenging datasets further demonstrates that the proposed outlier filter is robust to genomic heterogeneity, enabling accurate survival and hazard rate estimation even in the presence of multiple strains, alleles, or repetitive sequences.

## 3 Discussion

There remains room for further improvement in the skiver pipeline. In particular, performance on datasets with high error rates or low sequencing coverage could potentially be enhanced through adaptive selection of the parameters *k* and *v*. In addition, skiver can be fine-tuned to other types of data, such as amplicon sequencing (**Supplementary Table S11**). The error rate estimation results of skiver can also support a future generation of *k*-mer-based algorithms for alignment-free variant calling, read error correction, and other sequence analysis tasks.

Beyond applications to sequencing reads, the skiver framework could also be extended to collections of genomes. In this setting, the hazard rate can be interpreted as the probability that the next genomic position differs due to mutation. Under this formulation, skiver could enable efficient estimation of mutational spectra across large genome collections, providing a scalable approach for large-scale mutational spectra analysis.

## 4 Conclusion

In this study, we present skiver, a reference-free and alignment-free framework for profiling sequencing errors by using (*k, v*)-mer sketches. By modeling the time to first error as a discrete Weibull distribution, skiver provides direct estimates of survival and hazard rates, enabling a more expressive characterization of sequencing error processes than conventional single-number error metrics.

Across both simulated and real datasets, skiver consistently produces accurate survival and hazard rate estimates under a wide range of sequencing conditions. Skiver further enables reliable profiling of sequencing error spectra. The inferred substitution, insertion, and deletion patterns across Illumina, PacBio, and Nanopore datasets closely match previously reported platform-specific error characteristics. The proposed outlier filter substantially improves robustness in datasets containing multiple strains, alleles, or repetitive regions, enabling accurate estimation of error rates of reads coming from complex microbial communities and diploid genomes. Finally, skiver is computationally efficient, achieving low runtime and memory usage, making it suitable as a lightweight pre-processing step in sequence analysis pipelines.

We anticipate that integrating our algorithm into diverse bioinformatics pipelines will enable more reliable identification of high- and low-quality sequencing runs while providing insights for downstream tasks such as variant calling and genome assembly. More broadly, the principles underlying our approach may inspire further algorithmic advances in variant calling, consensus sequence generation, and alignment-free methods for sequence error correction.

## 5 Methods

### 5.1 (*k, v*)-mer sketches

#### Definition 3

A (*k, v*)-mer (as shown in **Figure 6.A**) is a segment of DNA of length *k* + *v*, with the first *k* bases being the *key* and the last *v* bases being the *value*.

**Fig. 6.**
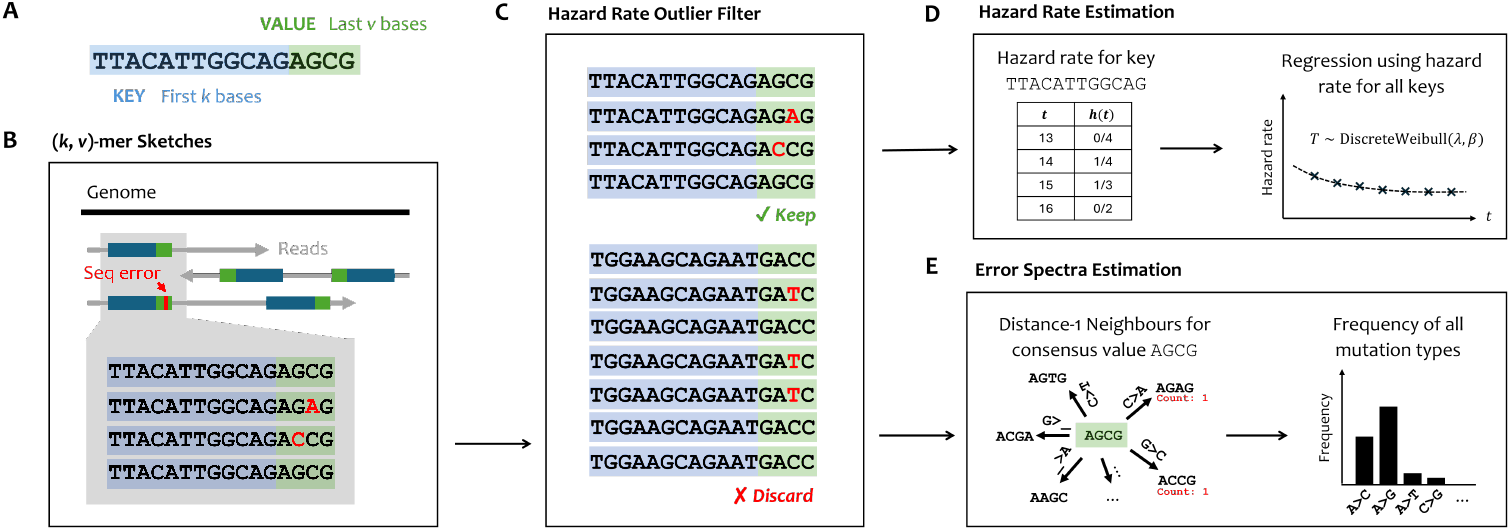
The workflow of skiver. **A**. The structure of a (*k, v*)-mer. **B**. Extracting (*k, v*)-mer sketches from the read set. (*k, v*)-mers with the same keys are gathered together to infer the consensus value and identify sequencing errors. **C**. The sketched (*k, v*)-mers go through a hazard rate outlier filter to exclude keys that are associated with multiple values with high counts. **D**. The hazard rate is calculated for each key and used to fit the parameters of a discrete Weibull distribution. **E**. We find all the distance-1 neighbors of the consensus and infer the frequency of each type of sequencing error (insertion, deletion, and substitution).

The structure of a (*k, v*)-mer is similar in spirit to the anchor-target *k*-mer pairs in SPLASH [38, 39] and strobemers [40], but different in that our key and value must be adjacent for accurate error profiling, whereas the anchor-target pair can be separated in the reads. In this paper, we focus on using this structure for sequencing error rate and spectra estimation.

A (*k, v*)-mer sketch of a set of sequenced reads is created with the following steps (**Figure 6.B**):

- **Step 1**. We extract all the (*k, v*)-mers from the reads and their reverse complement. Optionally, only the forward strand of the reads is used.
- **Step 2**. We subsample roughly 1*/c* (*k, v*)-mers with FracMinHash [41]. In particular, given a random hash function *H* : Σ^*k*^ →(0, 1), we subsample a (*k, v*)-mer if the hash value of its key is less than 1*/c*.
- **Step 3**. All the subsampled (*k, v*)-mers are stored in a hash table that maps a key to the set of associated values along with the number of times they appear in the read set. The most frequently appearing value is identified as the *consensus*.

Here, *k* is chosen to be large such that the keys are mostly unique in the set of sequenced genomes. The keys are used as positional identifiers in the genomes. We then identify variation in the associated values to determine possible sequencing errors or mutations. By default, both *k* and *v* are chosen to be 17, which generally works well across different real sequencing reads with various error rates (**Supplementary Figure S7**). *c* is chosen to be 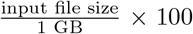 by default such that the number of sketched (*k, v*)-mers is around 10 million, enabling efficient processing on personal computers.

It is possible to show that given a per-base error rate of *ε*, when the number of times a key appears in the read set is *N*_key_ = Ω((1−*ε*)^−2*v*^ log *v*), the value with the highest count (consensus) matches the true value from the sequenced genome with high probability (**Supplementary Note S1**). In practice, we choose a default threshold *N*_key_ = 10, and only use keys above this coverage threshold for subsequent tasks, and show that this is sufficient for the evaluated datasets.

If a reference genome is provided, we also extract the set of (*k, v*)-mers from the genome using the same hash function. If a key *K* is associated with multiple different values in the reference genome, which indicates a repeat sequence, the key is discarded. Otherwise, the unique value associated with the key is regarded as the *consensus*. Since the consensus is obtained from the reference, a lower threshold for key multiplicity is set (*N*_key_ = 1) to enable profiling of lower coverage read sets. In this mode, it is recommended to use the full set of correct reference genomes, as incomplete reference sets can yield unstable error rate estimations (**Supplementary Figure S11**).

### 5.2 Estimating hazard rate and survival rate

In this work, we assume that *T* follows a discrete Weibull distribution with parameters *λ* and *β*, which is often used for discrete-time survival analysis. This assumption comes from the empirical observation that hazard rates *h*(*t*) in real sequencing datasets decrease with *t* (**Figure 4.A**), and the survival rate *S*(*t*) fits well to the curve *S*(*t*) = exp(−*λt*^*β*^) (**Supplementary Figure S9**). This model also produces better survival rate estimation than the constant hazard rate model, as we show in **section 2.6**.

Our model differs from existing error models which typically assume a constant error rate [15], essentially assuming *β* = 1. In real datasets, especially Nanopore and Illumina, the best fit parameter 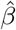 is much smaller than 1. This indicates a decreasing hazard rate, and is often interpreted as evidence for heterogeneity in the failure process [42], or heterogeneous sequencing error rate across reads in our case.

Given a (*k, v*)-mer sketch of the sequenced reads, we count *N*_*K,t*_, the number of (*k, v*)-mers that have key *K* and match with the consensus up to the *t*-th base. For example, in the example of **Figure 6.B**, 4 (*k, v*)-mers share the key *K* = TTACATTGGCAG. The consensus value is taken as the value with the highest count, in this case, `AGCG`. Two of the (*k, v*)-mers differ from the consensus in the 14th and 15th base, respectively. We therefore have *N*_*K*,12_ = *N*_*K*,13_ = 4, *N*_*K*,14_ = 3, *N*_*K*,15_ = *N*_*K*,16_ = 2. This process is repeated for all keys. We take the maximum likelihood estimate to estimate the hazard rate,

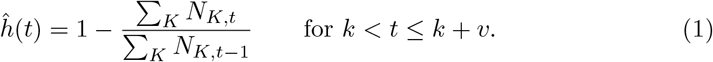

This allows us to estimate *h*(*t*) in a small interval between *k* and *k* + *v*. Under the assumption that *T* follows a discrete Weibull distribution, we have for large *t* (**Supplementary Note S2**),

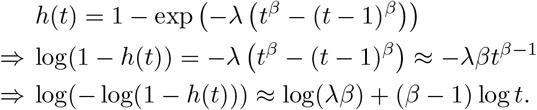

This transformation (complementary log-log) is widely used in discrete survival analysis [43]. We can then perform a ridge regression with Huber loss of log(− log(1−*h*(*t*))) vs. log *t* in the range [*k* + 1, *k* + *v*]. Let *a* and *b* be the estimated slope and intercept. Then, we have

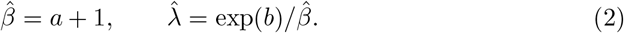

The survival rate can then be estimated by directly plugging the estimated parameters, 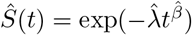, and the sequencing error rate is estimated to be 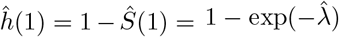. This process takes time and space that are linear in *v* and the number of keys in the sketch.

### 5.3 Estimating error spectra

To estimate the composition of errors (types of substitutions, insertions, and deletions), we make an additional assumption that the error composition is roughly constant and is independent of *t*. In other words, for a given single base error type *e* (such as substitution `C→A`), the hazard rate of this type of error occurring can be expressed as

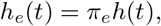

where *π*_*e*_ is a constant value. The probability of the type of error *e* happening exactly once in the value, while no other error is present, is

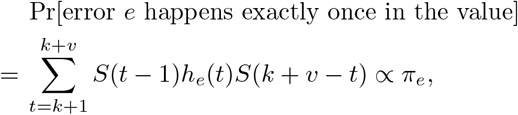

given that *k* and *v* are fixed. This assumption is based on the observation that the proportions of each error type happening exactly once at position *t* are roughly constant across *t* (**Supplementary Figure S10**). Under this assumption, we can estimate *π*_*e*_ by the frequency of *e* happening exactly once in the value.

After identifying the consensus value for each key, we find the set of all pairs (`value, edit type`), where the value can be obtained from the consensus via one edit of `edit type` (such as `A →C`). An example of the neighbor set can be found in **Supplementary Table S12**. If a neighbor can be reached via multiple types of edit, we mark the `edit type` as `Ambiguous`.

We then count the number of times each distance-1 neighbor appears in the (*k, v*)-mer sketch. In the example in **Figure 6.E**, the neighbors corresponding to the single base substitutions `C →A` and `G →C` appear once respectively. This process is repeated for all keys, which gives us the total number of times each`edit type` has appeared. The frequency of an `edit type` is estimated to be its count divided by the sum of counts of all the profiled error types. The `Ambiguous` neighbors are not counted.

There are at most 11*v* distance-1 neighbors given a consensus value. As a result, the time needed to count the error frequencies is only *O* ((*v·*#keys) +#values), where #keys and #values are the total number of keys and values in the (*k, v*)-mer sketch. This is much more efficient than the case in which we align all values to the consensus, which can take time *O*(*v*^2^ *·* #values).

### 5.4 Estimating the dependence of error rate on covariates

We can also estimate the conditional hazard rates *h*(*t* |*X*_*i*_) assuming a Cox proportional hazard rate model with the complementary-log-log link [43],

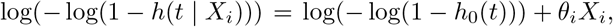

where *X*_*i*_ represents the covariates such as Phred quality scores, GC-content of the read, or the pore/well from which the read was generated. We choose the baseline

#### Algorithm 1

Outlier filter in hazard rate estimation

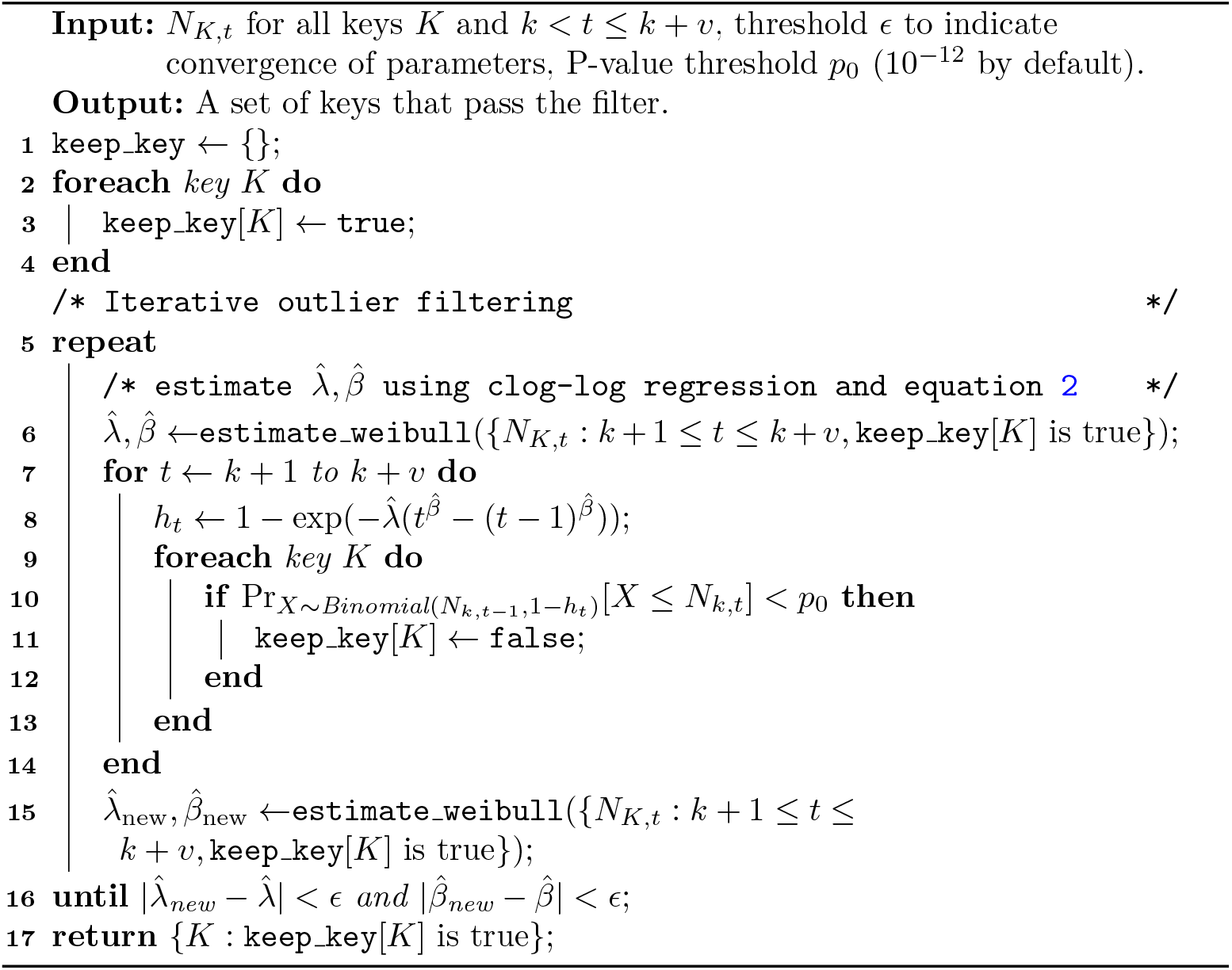

hazard *h*_0_(*t*) to be the hazard rate 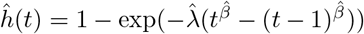 estimated using all the data points. We then calculate 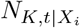 to be the number of bases in key *K* that satisfies *X*_*i*_ and agrees with the consensus up to the *t*-th base, and estimate

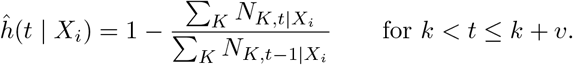

The individual scale parameters *θ*_*i*_ are then estimated using the mean of the difference between 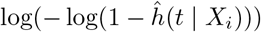and 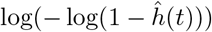.

### 5.5 Dealing with repeats, multiple alleles or strains

In real datasets, it is common that the sequenced genomes contain long repetitive regions and heterozygous sites in the case of the human genome, or multiple coexisting strains of the same species in the case of metagenomic samples. In these cases, a key can be associated with multiple values with high counts in the reads, causing overestimation of the hazard rate (**Figure 2.E**).

Suppose the hazard rate at the *t*-th base is *h*(*t*), the number of (*k, v*)-mers with key *K* that agree with the consensus up to the (*t* − 1)-th base is *N*_*K,t*−1_. Assuming independence across values, each of these *N*_*K,t*−1_ values continue to agree at the *t*-th base with probability 1 − *h*(*t*). Therefore,

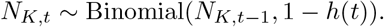

A key observation is that if a key *K* is associated with multiple values that have high counts, *N*_*K,t*_ is going to be significantly smaller for some *t* ∈ [*k* + 1, *k* + *v*]. We therefore use a simple outlier filter as shown in **Algorithm 1** that filters all *K* that have a significantly smaller *N*_*K,t*_, assuming that *N*_*K,t*_ ~Binomial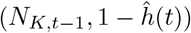.

If the reference genome is provided, this filter is disabled by default, as keys that are associated with multiple values are already discarded.

## Declarations

### Ethics approval and consent to participate

Not applicable.

### Consent for publication

Not applicable.

### Availability of data and materials

The implementation of skiver is available on GitHub at https://github.com/GZHoffie/skiver [44] and on Zenodo at 10.5281/zenodo.21817970 [45] under the MIT License. The datasets generated during and/or analysed during the current study, as well as the code for reproducibility, are available on GitHub at https://github.com/GZHoffie/skiver-test.

### Authors’ contributions

Conceptualization, Z.G. and N.N.; methodology, Z.G. and N.N.; software, Z.G.; data curation, Z.G.; investigation, Z.G.; writing-–original draft, Z.G. and P.S.; writing-–review & editing, Z.G., P.S., L.W., and N.N.; supervision, L.W. and N.N.

## Acknowledgments

Not applicable.

