## Supplementary Materials for "Skiver: reference-free quality control of metagenomic sequencing datasets using (*k, v*)-mer sketches"

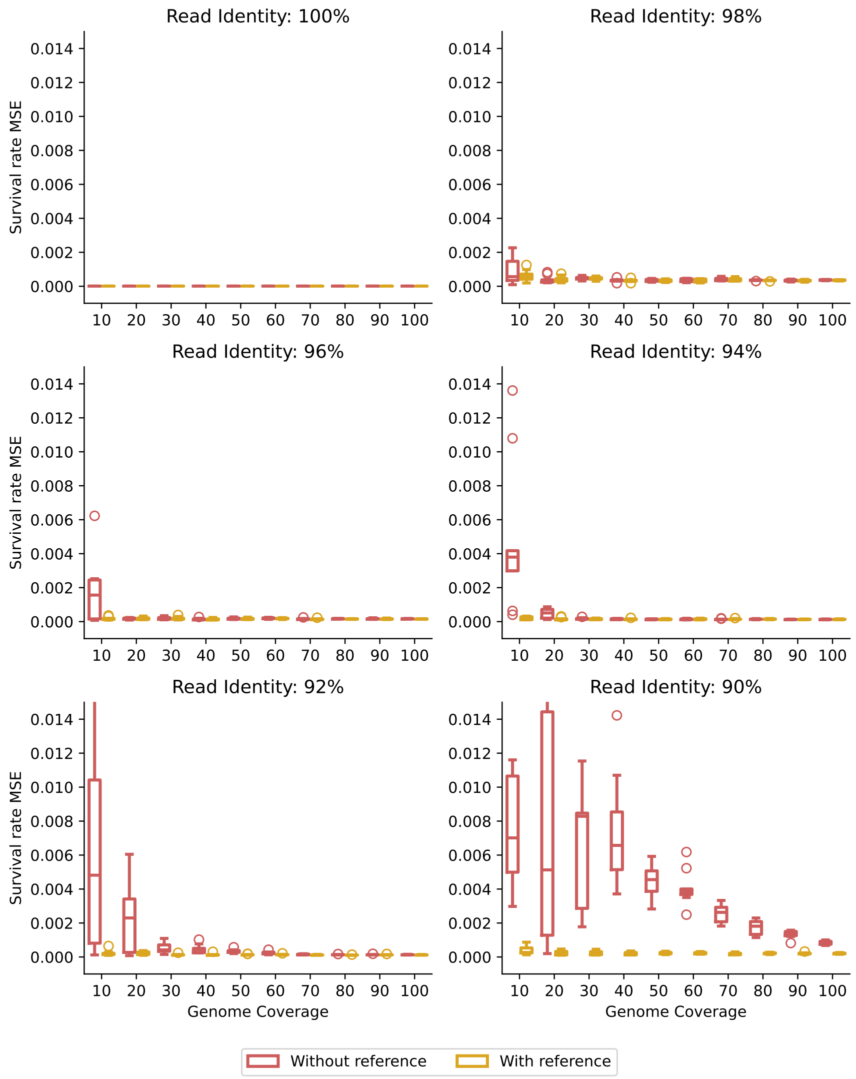


Figure S1: $MSE_S$ of skiver using Badread's simulated data.

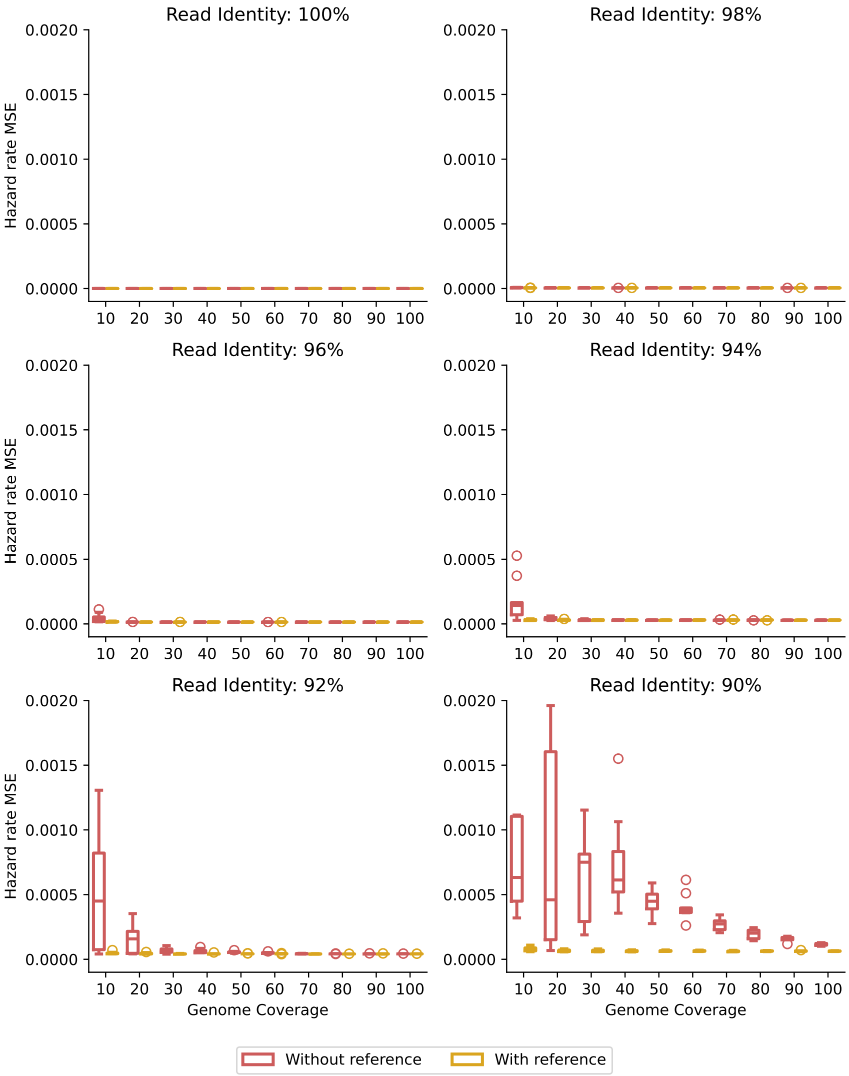


Figure S2: $\text{MSE}_h$ of skiver using Badread's simulated data.

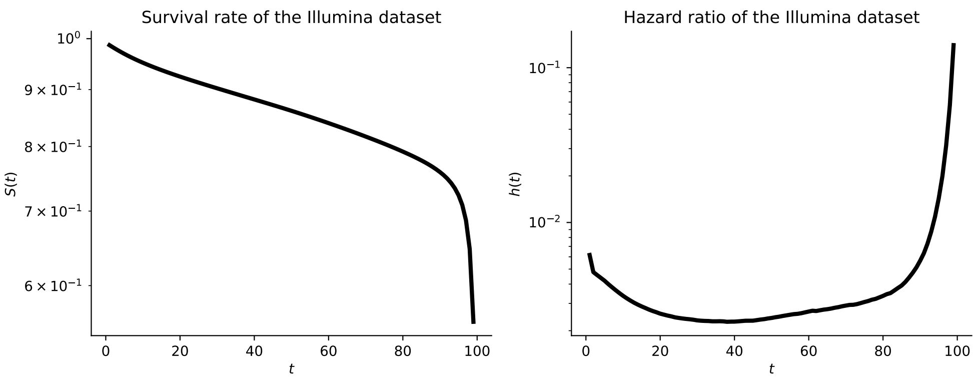


Figure S3: The observed survival rate and hazard rate of the *B. subtilis, Illumina* dataset. This dataset has a read length of 100. Due to the property of Illumina reads that the probability of having an error at the ends of the read is higher, this dataset exhibits a decreasing hazard rate for small $t$ and an increasing hazard rate for large $t$, showing a "bathtub" curve in the hazard rate plot.

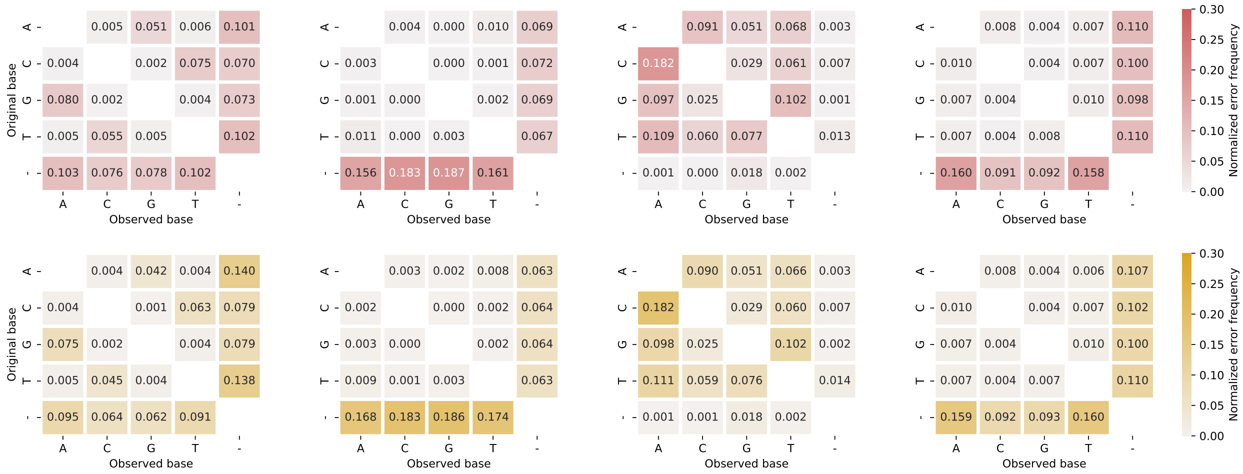


Figure S4: Estimated sequencing error spectra with skiver on metagenomic datasets, with (top) and without (bottom) reference genome.

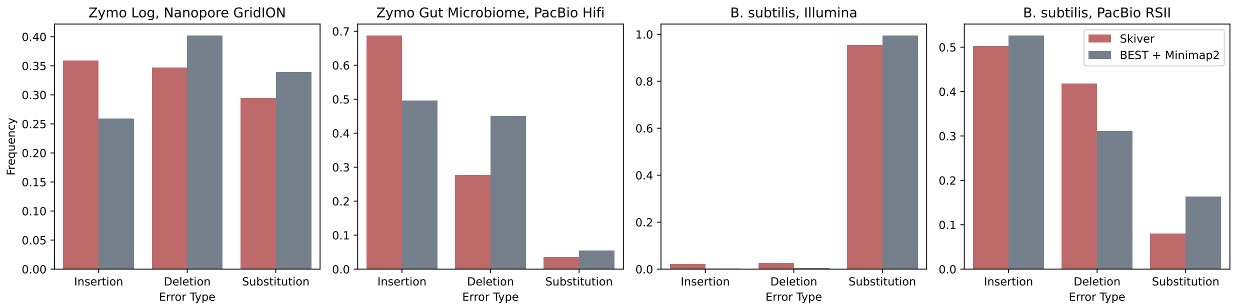


Figure S5: Estimated error type composition of Minimap2 + BEST and skiver.

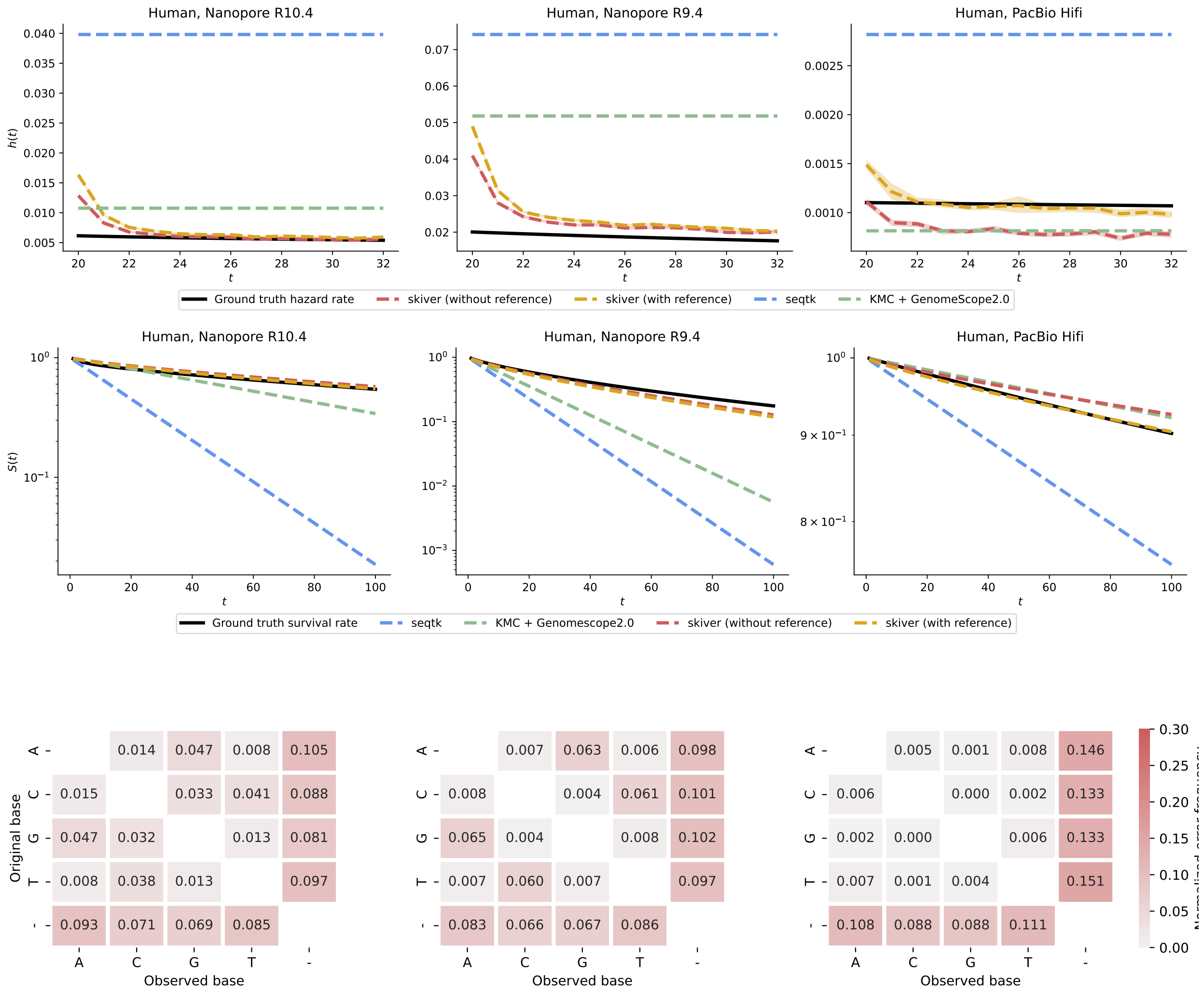


Figure S6: Estimated hazard rate (top), survival rate (middle) and error spectrum (bottom) of the HG002 sequenced reads on different platforms.

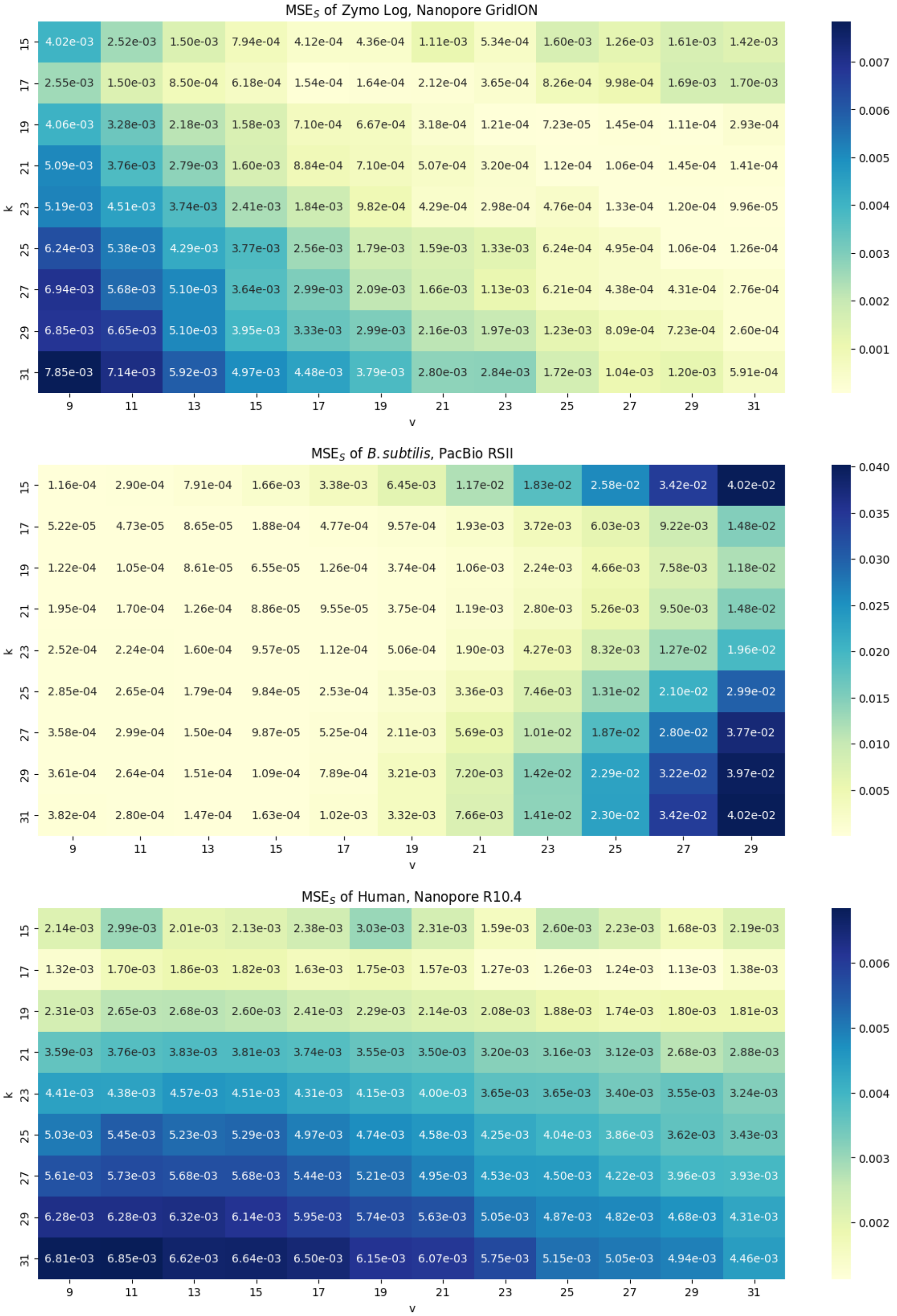


Figure S7: MSE$_S$ of skiver using three real datasets using various combinations of $k$ and $v$. In general, MSE$_S$ doesn't vary a lot as $k$ and $v$ change. For reads with a low error rate, a larger $v$ would be preferable; and for high error rate, a smaller $v$ is preferable. $k = v = 17$ is a good choice across all datasets.

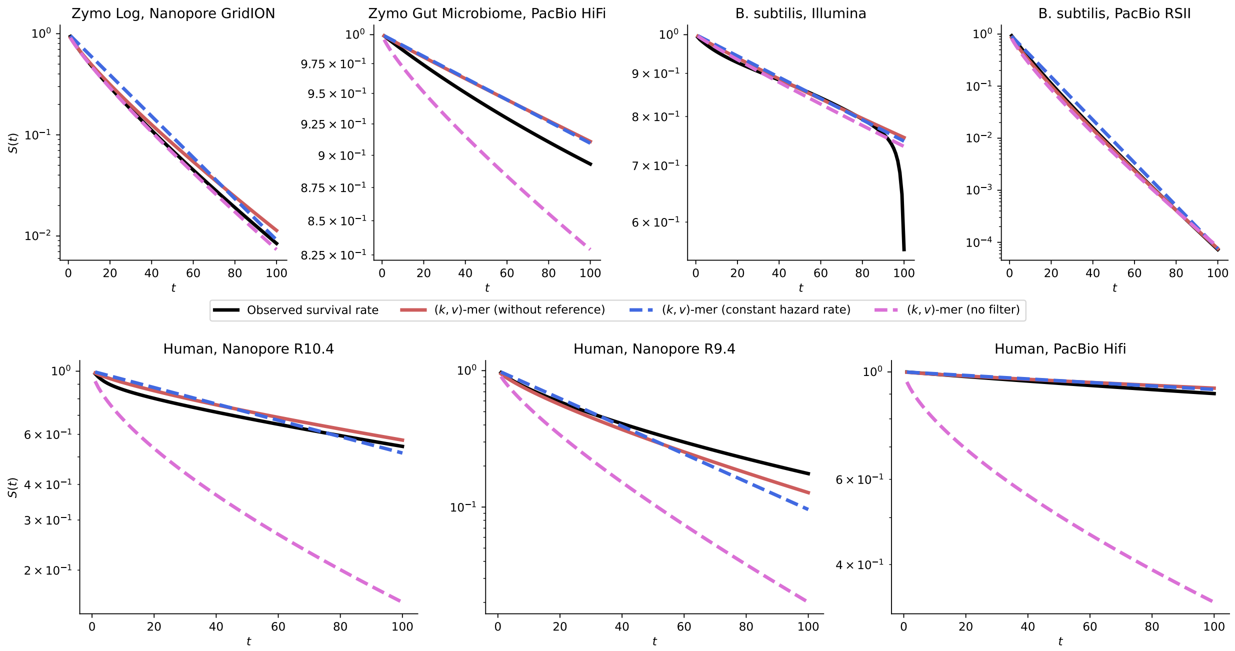


Figure S8: Ablation study. The predicted survival rate of default skiver (red lines), skiver assuming a constant hazard rate model instead of a Weibull distribution (blue lines), skiver without filtering the hazard rate outliers (pink lines).

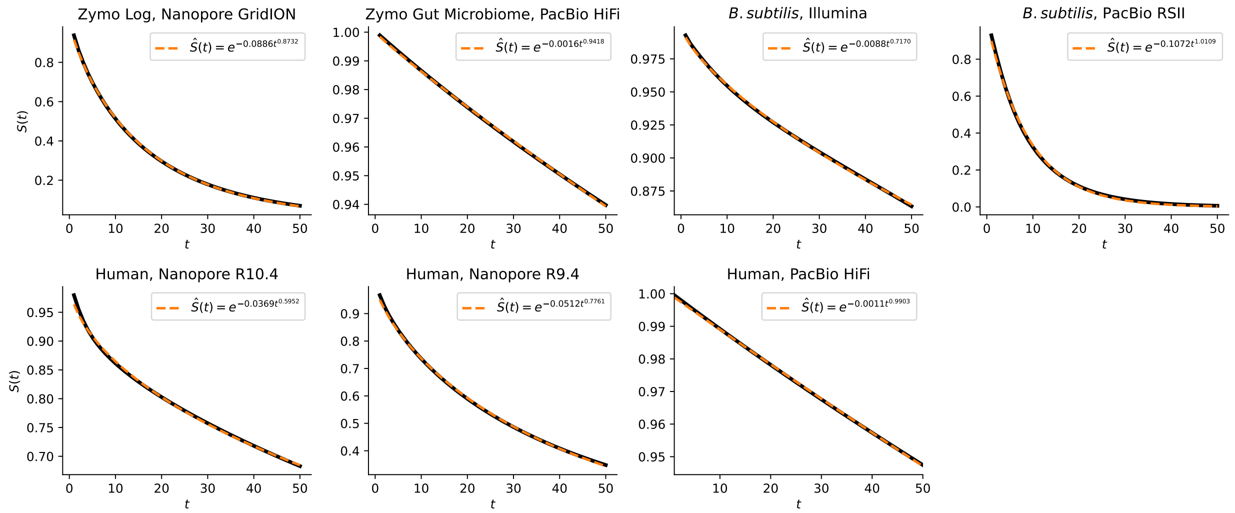


Figure S9: Least-square best fits of the observed survival rate of various real sequencing datasets, assuming $T$ follows Weibull distribution, the survival rate is best expressed as $S(t) = \exp(-\lambda t^{\beta})$. We have $\beta < 1$ in most of the best fits, which indicates a heterogeneous error rate among the reads.

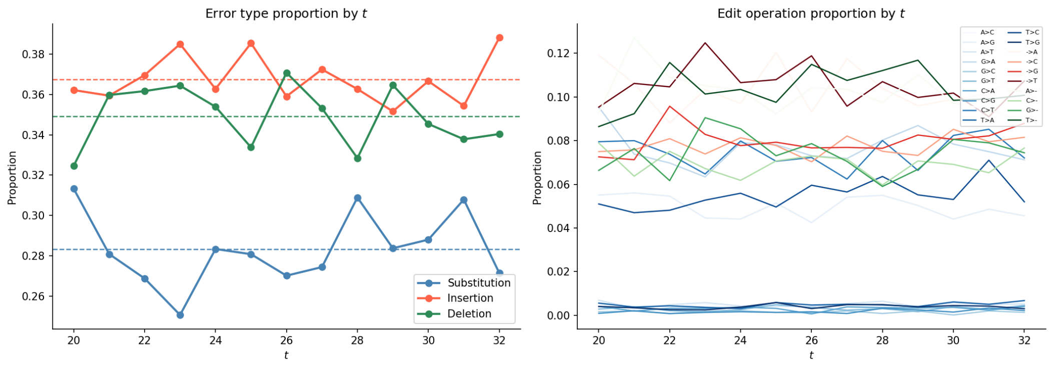


(a) Zymo Log, Nanopore GridION

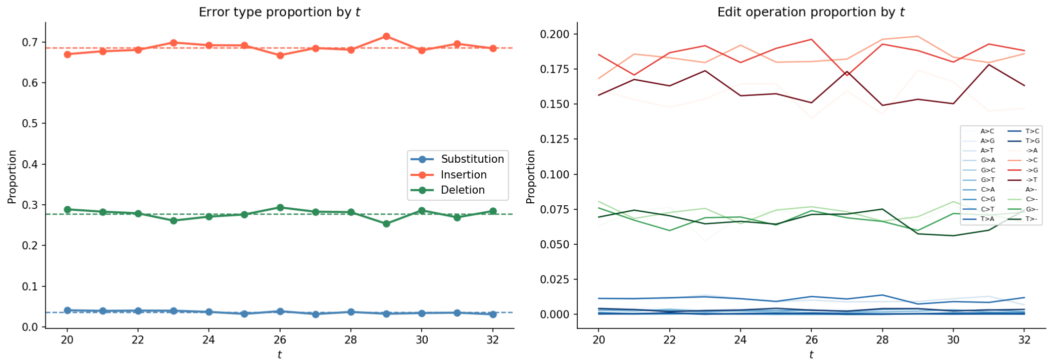


(b) Zymo Gut Microbiome, PacBio Hifi

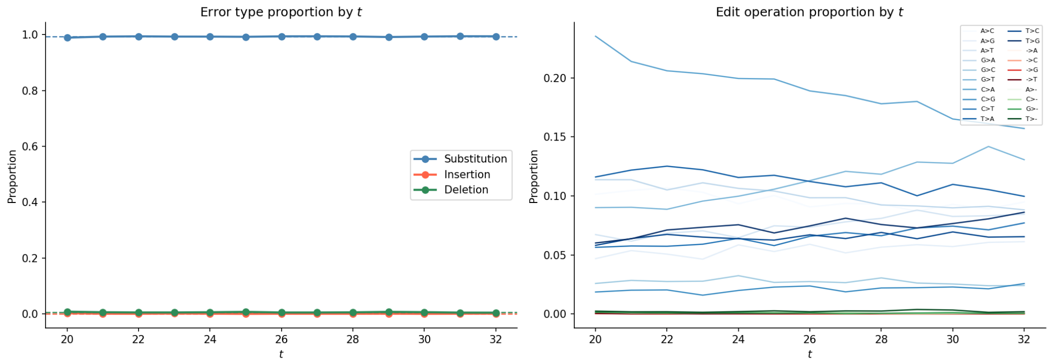


(c) *B. subtilis*, Illumina

Figure S10: Dependence of proportion of each type of error (insertion, deletion, substitution) on $t$.

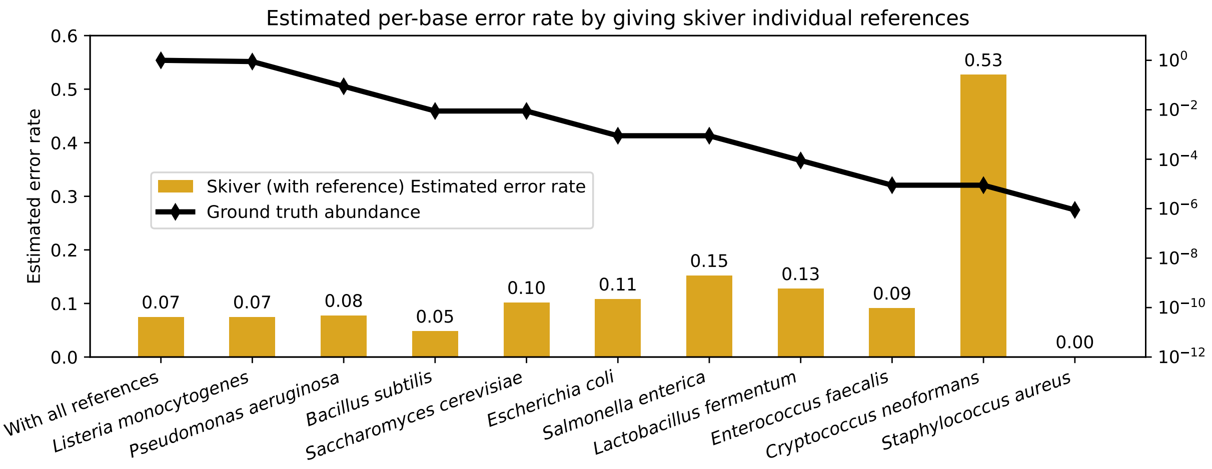


Figure S11: Per-base error rate estimate of skiver on the *Zymo Log, Nanopore GridION* dataset using individual genome as reference. The estimates of skiver (with reference) show a small variance when using different references, except for *Cryptococcus neoformans* and *Staphylococcus aureus*, where the coverage of the genomes is too small to make a reasonable estimate.

Table S1: Estimation of skiver using the *B. subtilis, Illumina* dataset with 0, 5, 10 bases trimmed at the two ends of the reads.

| Trimmed bases | Per-base error rate estimation | $MSE_S$ | $MSE_h$ |
|---:|---:|---:|---:|
| 0 | 0.004018 | $6.8 \times 10^{-4}$ | $2.3317 \times 10^{-4}$ |
| 5 | 0.003571 | $7.9 \times 10^{-4}$ | $2.3330 \times 10^{-4}$ |
| 10 | 0.003392 | $7.38 \times 10^{-4}$ | $2.3285 \times 10^{-4}$ |

Table S2: $MSE_h$ for the benchmarked methods across real sequencing datasets. The lowest value for each dataset is shown in bold and the second-lowest value is underlined.

| Dataset | seqtk | KMC + GenomeScope2.0 | Skiver | Skiver (with reference) |
|---|---:|---:|---:|---:|
| Zymo Log, Nanopore GridION | $4.36 \times 10^{-3}$ | $1.44 \times 10^{-4}$ | $1.75 \times 10^{-5}$ | $\mathbf{1.13 \times 10^{-5}}$ |
| Zymo Gut Microbiome, PacBio Hifi | $1.79 \times 10^{-8}$ | $4.53 \times 10^{-5}$ | $5.08 \times 10^{-8}$ | $\mathbf{8.75 \times 10^{-9}}$ |
| *B. subtilis*, Illumina | $\mathbf{2.24 \times 10^{-4}}$ | $\mathbf{2.24 \times 10^{-4}}$ | $2.33 \times 10^{-4}$ | $2.32 \times 10^{-4}$ |
| *B. subtilis*, PacBio RSII | $7.63 \times 10^{-3}$ | $7.67 \times 10^{-4}$ | $7.04 \times 10^{-5}$ | $\mathbf{4.62 \times 10^{-5}}$ |
| Human, Nanopore R10.4 | $1.15 \times 10^{-3}$ | $3.41 \times 10^{-5}$ | $4.87 \times 10^{-6}$ | $\mathbf{3.85 \times 10^{-6}}$ |
| Human, Nanopore R9.4 | $3.27 \times 10^{-3}$ | $1.23 \times 10^{-3}$ | $\mathbf{1.36 \times 10^{-5}}$ | $2.00 \times 10^{-5}$ |
| Human, PacBio Hifi | $3.21 \times 10^{-6}$ | $5.38 \times 10^{-8}$ | $8.64 \times 10^{-8}$ | $\mathbf{3.60 \times 10^{-8}}$ |

Table S3: Estimated per-base error rates of the benchmarked tools. The profiled error rates from skiver are the estimated $\hat{h}(1) = 1 - \exp(-\hat{\lambda})$. For Minimap2 + BEST, the gap-compressed error rate is reported.

| Dataset | Minimap2 + BEST | seqtk | KMC + Genomescope2.0 | skiver | skiver (with reference) |
|---|---:|---:|---:|---:|---:|
| Zymo Log, Nanopore GridION | 0.089030 | 0.1122 | 0.000373 | 0.0887 | 0.0743 |
| Zymo Gut Microbiome, PacBio Hifi | 0.001441 | 0.0011 | 0.000079 | 0.0011 | 0.0015 |
| *B. subtilis*, Illumina | 0.007221 | 0.0049 | 0.000057 | 0.0040 | 0.0041 |
| *B. subtilis*, PacBio RSII | 0.103340 | 0.1778 | 0.001173 | 0.1503 | 0.1141 |
| Human, Nanopore R10.4 | 0.028530 | 0.0398 | 0.010760 | 0.0150 | 0.0172 |
| Human, Nanopore R9.4 | 0.047698 | 0.0741 | 0.051815 | 0.0517 | 0.0587 |
| Human, PacBio Hifi | 0.001175 | 0.0028 | 0.000814 | 0.0015 | 0.0021 |

Table S4: Estimation of the percentage of unclassified reads of the metagenomic samples. We use Minimap2 to map the reads from the Zymo mock communities and the Zymo reference database to find the percentage of reads not mapped as the ground truth.

| Dataset | Minimap2 (ground truth) | Default sylph | Skiver + sylph |
|---|---:|---:|---:|
| Zymo Log, Nanopore GridION | 89.9126% | 36.3728% | 89.4377% |
| Zymo Gut Microbiome, PacBio Hifi | 96.9265% | 95.5556% | 96.9850% |

Table S5: Per-base error rate (Insertion/Deletion/Substitution rate) estimation of skiver on real metagenomic datasets. Skiver without outlier filter is biased heavily upwards.

| Dataset | Skiver | Skiver (no filter) |
|---|---|---|
| Human Gut, Nanopore PromethION | 7.27% (2.099% / 2.900% / 2.271%) | 9.628% (2.711% / 3.934% / 2.983%) |
| Soil sample, Illumina NovaSeq | 0.448% (0.001% / 0.001% / 0.446%) | 6.565% (0.055% / 0.064% / 6.446%) |
| Water sample, Illumina NovaSeq | 0.462% (0.001% / 0.002% / 0.459%) | 5.289% (0.036% / 0.030% / 5.223%) |

Table S6: Estimated error rates of the variants of the default skiver algorithm: "constant hazard rate" - assuming that the hazard rate is constant; "no filter" - when no filter of outliers is applied. The profiled error rates are the estimated $\hat{h}(1) = 1 - \exp(-\hat{\lambda})$.

| Dataset | Default skiver | constant hazard rate | no filter |
|---|---|---|---|
| Zymo Log, Nanopore GridION | 0.088681 | 0.045869 | 0.089319 |
| Zymo Gut Microbiome, PacBio Hifi | 0.001083 | 0.000948 | 0.004329 |
| *B. subtilis*, Illumina | 0.004018 | 0.002898 | 0.004162 |
| *B. subtilis*, PacBio RSII | 0.150313 | 0.090512 | 0.181092 |
| Human, Nanopore R10.4 | 0.014999 | 0.006597 | 0.078046 |
| Human, Nanopore R9.4 | 0.051685 | 0.023164 | 0.095097 |
| Human, PacBio Hifi | 0.001487 | 0.000833 | 0.046505 |

Table S7: $\text{MSE}_h$ across different datasets for the variants of the Skiver algorithm.

| Dataset | Default skiver | constant hazard rate | no filter |
|---|---|---|---|
| Zymo Log, Nanopore GridION | $1.75 \times 10^{-5}$ | $5.80 \times 10^{-5}$ | $\mathbf{9.97 \times 10^{-6}}$ |
| Zymo Gut Microbiome, PacBio Hifi | $\mathbf{5.08 \times 10^{-8}}$ | $5.09 \times 10^{-8}$ | $6.66 \times 10^{-7}$ |
| *B. subtilis*, Illumina | $2.33 \times 10^{-4}$ | $\mathbf{2.32 \times 10^{-4}}$ | $\mathbf{2.32 \times 10^{-4}}$ |
| *B. subtilis*, PacBio RSII | $7.04 \times 10^{-5}$ | $\mathbf{7.02 \times 10^{-5}}$ | $1.61 \times 10^{-4}$ |
| Human, Nanopore R10.4 | $\mathbf{4.87 \times 10^{-6}}$ | $1.23 \times 10^{-5}$ | $1.86 \times 10^{-4}$ |
| Human, Nanopore R9.4 | $\mathbf{1.36 \times 10^{-5}}$ | $6.49 \times 10^{-5}$ | $4.67 \times 10^{-4}$ |
| Human, PacBio Hifi | $8.64 \times 10^{-8}$ | $\mathbf{4.63 \times 10^{-8}}$ | $1.23 \times 10^{-4}$ |

Table S8: Running time (in seconds) of the tools on different datasets.

| Dataset | KMC + Genomescope2.0 | seqtk | Minimap2 + BEST | Skiver |
|---|---|---|---|---|
| Zymo Log, Nanopore GridION | 934.24 | 310.20 | 3591.82 | 322.45 |
| Zymo Gut Microbiome, PacBio Hifi | 1164.23 | 221.79 | 4753.00 | 349.93 |
| *B. subtilis*, Illumina | 304.16 | 66.82 | 1078.41 | 93.11 |
| *B. subtilis*, PacBio RSII | 73.11 | 12.12 | 441.58 | 99.99 |
| Human, Nanopore R10.4 | 4631.00 | 1823.54 | 29867.00 | 1078.43 |
| Human, Nanopore R9.4 | 3858.00 | 1679.65 | 5942.00 | 1103.88 |
| Human, PacBio Hifi | 6331.00 | 1627.24 | 20533.00 | 1253.03 |

Table S9: Memory (in GB) of the tools on different datasets.

| Dataset | KMC + Genomescope2.0 | seqtk | Minimap2 + BEST | Skiver |
|---|---|---|---|---|
| Zymo Log, Nanopore GridION | 29.81 | 0.20 | 3.86 | 2.25 |
| Zymo Gut Microbiome, PacBio Hifi | 29.75 | 0.05 | 3.15 | 5.51 |
| *B. subtilis*, Illumina | 23.29 | 0.002 | 0.87 | 0.65 |
| *B. subtilis*, PacBio RSII | 11.91 | 0.10 | 3.62 | 4.64 |
| Human, Nanopore R10.4 | 29.89 | 1.55 | 22.02 | 3.27 |
| Human, Nanopore R9.4 | 29.96 | 3.10 | 22.02 | 3.25 |
| Human, PacBio Hifi | 29.93 | 0.05 | 22.02 | 5.86 |

Table S10: Size of the $(k, v)$-mer sketches and the time needed to construct the sketches.

| Target | Platform | # of bases | Size of sketch (MBytes) | Time to sketch (s) |
|---|---|---|---|---|
| Zymo Log | Nanopore GridION | 16.5 Gbp | 237 | 288.42 |
| Zymo Gut Microbiome | PacBio HiFi | 18 Gbp | 360 | 285.49 |
| *Bacillus subtilis* | Illumina | 4.7 Gbp | 196 | 80.89 |
| *Bacillus subtilis* | PacBio RSII | 1.6 Gbp | 618 | 40.72 |
| HG002 | Nanopore R10.4 | 60.6 Gbp | 224 | 1006.11 |
| | Nanopore R9.4 | 64.2 Gbp | 280 | 1037.65 |
| | PacBio HiFi | 109.06 Gbp | 434 | 1173.74 |

Table S11: Skiver's results on some amplicon data.

| Target | Synthetic Bacterial Mix | Zymo_ontR9.4.1 |
|---|---|---|
| Platform | 16S Illumina MiSeq | 16S Nanopore PromethION |
| Source | NCBI: SRR1204979 | NCBI: SRR23176493 |
| Estimated per-base error rate | 0.00249 | 0.049791 |
| Sub./Ins./Del. rates | 0.246%/0.000%/0.003% | 2.317%/1.227%/1.435% |

Table S12: The full one-edit neighbor set of value AACG. Here, A $\to$ T denotes a substitution of a base A to T, $- \to$ A denotes an insertion of a base A, and A $\to -$ denotes a deletion of a base A. Note that when a base is deleted, a new base is filled in at the end to maintain the length of the value $v$. Thus, there are at most $4v$ possible neighbors for the delete operation. *Ambiguous* means that the neighbor can be reached using multiple edit operations. For example, AAGC can be obtained from AACG either by deleting the third base C and appending C, or by inserting G before the third position and truncating the final base. Therefore, this neighbor is labeled as ambiguous.

| Neighbor | Edit Operation | Neighbor | Edit Operation |
|---|---|---|---|
| AAAC | $- \to$ A | ATCG | A $\to$ T |
| TACG | A $\to$ T | ACGG | A $\to -$ |
| AGCG | A $\to$ G | CAAC | $- \to$ C |
| ACAC | $- \to$ C | TAAC | $- \to$ T |
| AAGA | C $\to -$ | ACGT | A $\to -$ |
| ATAC | $- \to$ T | AACT | *Ambiguous* |
| AACA | *Ambiguous* | AGAC | $- \to$ G |
| AAGC | *Ambiguous* | ACGC | A $\to -$ |
| GAAC | $- \to$ G | AAAG | C $\to$ A |
| ACGA | A $\to -$ | AATC | $- \to$ T |
| CACG | A $\to$ C | AAGG | *Ambiguous* |
| ACCG | A $\to$ C | AAGT | C $\to -$ |
| GACG | A $\to$ G | AATG | C $\to$ T |
| AACC | *Ambiguous* | | |

#### Supplementary Note S1

In this supplementary note, we give an informal theoretical lower bound for the coverage of the keys needed so that in the $(k, v)$-mer sketch, the value with the highest count (the *consensus* value) is the correct value in the sequenced genome with high probability.

For simplicity, we make the following assumptions:

1. each base in the reads has an independent and equal chance of being erroneous with probability $\varepsilon$.
2. we restrict attention to values with at most one sequencing error, and assume that the probability that a value with two or more errors is the most frequent erroneous value is negligible.

For a given key $K$, let $V^*$ denote the true value after the key in the underlying reference genome. Suppose the key $K$ appears $N$ times in the read set. Let $V_1, ..., V_N$ be the observed values associated with the key. The probability of the observed value agreeing with the ground truth $V^*$ is

$$p := \Pr[V_j = V^*] = (1 - \varepsilon)^v,$$

According to our first assumption.

Fix a particular distance-1 neighbor $w$ of $V^*$, which can be obtained from $V^*$ using one edit (one error among the $v$ bases), the probability of observing this neighbor in the read set is

$$q := \Pr[V_j = w] \le (1 - \varepsilon)^{v-1}\varepsilon$$

Let $C^*$ and $C_w$ denote the number of times $V^*$ and $w$ appears in the read set,

$$C^* := \sum_{j=1}^{N} I(V_j = V^*), \qquad C_w = \sum_{j=1}^{N} I(V_j = w),$$

where $I(\cdot)$ is an indicator function that is 1 if the statement inside is evaluated to true, and 0 otherwise. Our goal is to show that $C^* > C_w$ for all $w$ with high probability.

Let $D_j := I(V_j = V^*) - I(V_j = w)$, then each $D_1, \ldots, D_N$ are i.i.d. random variables that take value in $\{-1, 0, 1\}$. Let $S_N := \sum_{j=1}^{N} D_j$, we have

$$\mathbf{E}[D_j] = p - q, \qquad \mathbf{E}[S_N] = \mathbf{E}\left[\sum_{i=1}^{N} D_j\right] = N(p-q).$$

By Hoeffding's inequality, the probability of the count of $w$ exceeding the count of $V^*$ is

$$\begin{aligned}
\Pr[C^* \leq C_w] &= \Pr\left[S_N \leq 0\right] \\
&= \Pr\left[S_N - \mathbf{E}\left[S_N\right] \leq -N(p-q)\right] \\
&\leq \exp\left(-\frac{2N^2(p-q)^2}{4N}\right) = \exp\left(-\frac{N(p-q)^2}{2}\right).
\end{aligned}$$

There are at most $11v$ distance-1 neighbors of $V^*$. If the multiplicity of the key is sufficiently large, i.e.

$$N \geq \frac{2\log\frac{11v}{0.01}}{(p-q)^2} = \frac{2\log(1100v)}{\left((1-\varepsilon)^{v-1}(1-2\varepsilon)\right)^2}, \tag{1}$$

by union bound,

$$\begin{aligned}
\Pr[C^* > C_w \text{ for all } w] &\geq 1 - 11v \cdot \Pr[C^* > C_w] \\
&= 1 - 11v \cdot \exp\left(-\frac{N(p-q)^2}{2}\right) \\
&\geq 1 - 11v \cdot \frac{0.01}{11v} = 0.99,
\end{aligned}$$

This means that if the multiplicity of keys $N = \Omega((1-\varepsilon)^{-2v}\log v)$ then we have $C^* > C_w$ for all $w$ with high probability. The higher the coverage $N$ is, the higher the probability that $C^*$ is the highest count.

### Supplementary Note S2

In this supplementary note, we give a mathematical justification of the approximation of the relationship between the complementary log-log of the hazard rate and $t$,

$$\begin{aligned}
& h(t) = 1 - \exp\left(-\lambda\left(t^\beta - (t-1)^\beta\right)\right) \\
\Rightarrow\quad & \log(1 - h(t)) = -\lambda\left(t^\beta - (t-1)^\beta\right) \\
\Rightarrow\quad & \log\left(-\log(1-h(t))\right) = \log\lambda + \log\left(t^\beta - (t-1)^\beta\right)
\end{aligned}$$

Using a Taylor expansion, we have

$$(t-1)^\beta = t^\beta \left(1 - \frac{1}{t}\right)^\beta = t^\beta \left(1 - \frac{\beta}{t} + \frac{\beta(\beta-1)}{2t^2} + \mathcal{O}(t^{-3})\right),$$

so

$$\begin{aligned} t^\beta - (t-1)^\beta &= t^\beta \left(\frac{\beta}{t} - \frac{\beta(\beta-1)}{2t^2} + \mathcal{O}(t^{-3})\right) \\ &= \beta t^{\beta-1} \left(1 - \frac{\beta-1}{2t} + \mathcal{O}(t^{-2})\right), \end{aligned}$$

and therefore

$$\begin{aligned} \log\left(-\log(1-h(t))\right) &= \log\lambda + \log\left(t^\beta - (t-1)^\beta\right) \\ &= \log\lambda\beta + (\beta-1)\log t + \log\left(1 - \frac{\beta-1}{2t} + \mathcal{O}(t^{-2})\right) \\ &\approx \log\lambda\beta + (\beta-1)\log t. \end{aligned}$$

This approximation is based on the observation that $\beta \approx 1$ in the real dataset (**Supplementary Figure S9**) and $t \in [k+1, k+v]$ is large in the estimated hazard rates $h(t)$. It should be noted that this approximation is biased, but causes negligible error in the estimation of $\lambda$ and $\beta$.
